# Branch-specific clustered parallel fiber input controls dendritic computation in Purkinje cells

**DOI:** 10.1101/2024.01.25.577146

**Authors:** Gabriela Cirtala, Erik De Schutter

**Affiliations:** Computational Neuroscience Unit, Okinawa Institute of Science and Technology Graduate University, Okinawa, Japan

**Keywords:** Purkinje cell, heterogenous model, dendritic branches, dendritic spikes, bimodal response, ion channel conductance

## Abstract

Most central neurons have intricately branched dendritic trees that integrate massive numbers of synaptic inputs. Intrinsic active mechanisms in dendrites can be heterogenous and be modulated in a branch-specific way. However, it remains poorly understood how heterogenous intrinsic properties contribute to processing of synaptic input. We propose the first computational model of the cerebellar Purkinje cell with dendritic heterogeneity, in which each branch is an individual unit and is characterized by its own set of ion channel conductance densities. When simultaneously activating a cluster of parallel fiber synapses, we measure the peak amplitude of a response (PAR) and observe how changes in P-type calcium channel conductance density shift PAR from a linear one to a bimodal one including dendritic calcium spikes and vice-versa. These changes relate to the morphology of each branch. We show how dendritic calcium spikes propagate and how Kv4 channels block spreading depolarization to nearby branches.

## 1. Introduction

Cerebellar Purkinje cells (PCs) represent the sole output of the cerebellar cortex and are involved with encoding sensory and motor information. The extensive dendritic branching grants PCs with a unique architecture and allows them to process massive amount of information with great accuracy. PCs receive excitatory synaptic input from approximately 150,000 parallel fibers (PFs),^1^ which when activated, trigger local dendritic calcium spikes that are essential for inducing synaptic plasticity. ^2,3^

The active properties of PC dendrites were first proposed five decades ago by Llinás et al.^4^, and ever since, the scientific community has successfully uncovered many of their properties by combining modeling and experimental studies.^5–10^ An extensive review of how PC models were first developed and their immense contribution to the understanding of the active electrical properties in the dendrites of the central nervous system was done by Bower.^11^ It is well known that PC dendrites possess many different ion channels^12–17^ such as voltage dependent potassium channels (Kv4, Kv3), large conductance calcium-activated potassium channels (BK), small conductance calcium activated potassium channels (SK), high threshold P-type Calcium channels (CaP), etc.

While PC dendritic spikes can also be generated via climbing fiber activation,^5,18–20^ in this work we will focus on the far less studied dendritic calcium spikes triggered by strong local clustered parallel fiber (PF) activation.^2,3^ During the last decade, local computation in dendrites has been captured by multiple experimental studies that highlight the importance of branch-specific generated dendritic spikes on synaptic plasticity and information storage.^21–25^ Branco et al.^23^ discuss in detail dendritic branches acting as individual processing units, reviewing evidence of electrical, chemical, and translational compartmentalization on single branch scale, while showcasing that each branch may possess unique features given by the functional properties of its synaptic inputs. Not limited to the cerebellum, such branch-specific activity has been demonstrated in different areas of the brain like cortex^24,25^ and hippocampus.^25,26^

In the cerebellum, Zang and De Schutter^27^ proposed a computational model that unveiled the first evidence of localized PF dendritic spikes in a single neuron.^27^ The authors simulated clustered PF input by randomly distributing PF synapses on 22 manually defined branches. When examining their PAR with respect to increasing number of PF synapses they found that four of the branches exhibited a bimodal linear-step-plateau response, characterized by a linear increase of excitatory postsynaptic potential (EPSP) amplitude until a certain threshold, followed by a voltage jump caused by a dendritic calcium spike of about constant amplitude, while most of the branches had linear responses. These interesting results suggest that single PCs are capable of implementing their own branch specific multiplexed coding.^27,28^ Moreover, this raises many exciting scientific questions such as why is the response different between branches, how does each branch-characteristic morphology affect the PAR and how does this influence cerebellar coding and learning capacity?

The well-validated model proposed by Zang and De Schutter,^27^ like others previously developed,^8–10,27,29–32^ assumes dendritic ion channel conductance densities to be uniform throughout the PC spiny dendrite. This commonly employed assumption is traditionally made for the sake of simplicity, as considering different ion channel conductance densities would significantly increase the number of parameters used in the model and therefore the time required to properly tune these parameters. However, this common assumption may need to be reconsidered due to increasing evidence of ion channel heterogeneity across different dendrites, found in many different experimental studies in pyramidal cells^21,22^ and Purkinje cells.^2,33–36^

In this article we propose, to our knowledge, the first heterogeneous ion channel density model in which each branch of the PC is characterized by its own set of ion channel conductance densities. We show how modifying the biophysical properties of each branch modify its PAR, producing a shift in the synaptic gain curves from linear to bimodal linear-step-plateau and vice versa. We also discuss propagation of dendritic calcium spikes within PCs, and we propose a mechanism for blocking their progression onto nearby dendrites. Additionally, we discuss how co-activation of different dendritic branches changes the gain response and dendritic spike propagation.

## 2. Results

In this study, we continue the work of Zang and De Schutter^27^ to further explore the multiplexed coding strategies that PCs use in response to a clustered PF input.^2,3^ We split the dendritic tree in 22 different branches^27^ (**Figure 1A**), and we uniformly distribute PF synapses within each branch. Unlike previous literature, our model considers heterogenous ion channel densities for each branch in the dendritic tree. In this section, we show how altering biophysical properties shifts the PAR from linear to bimodal linear-step-plateau and vice versa.

**Figure 1:**
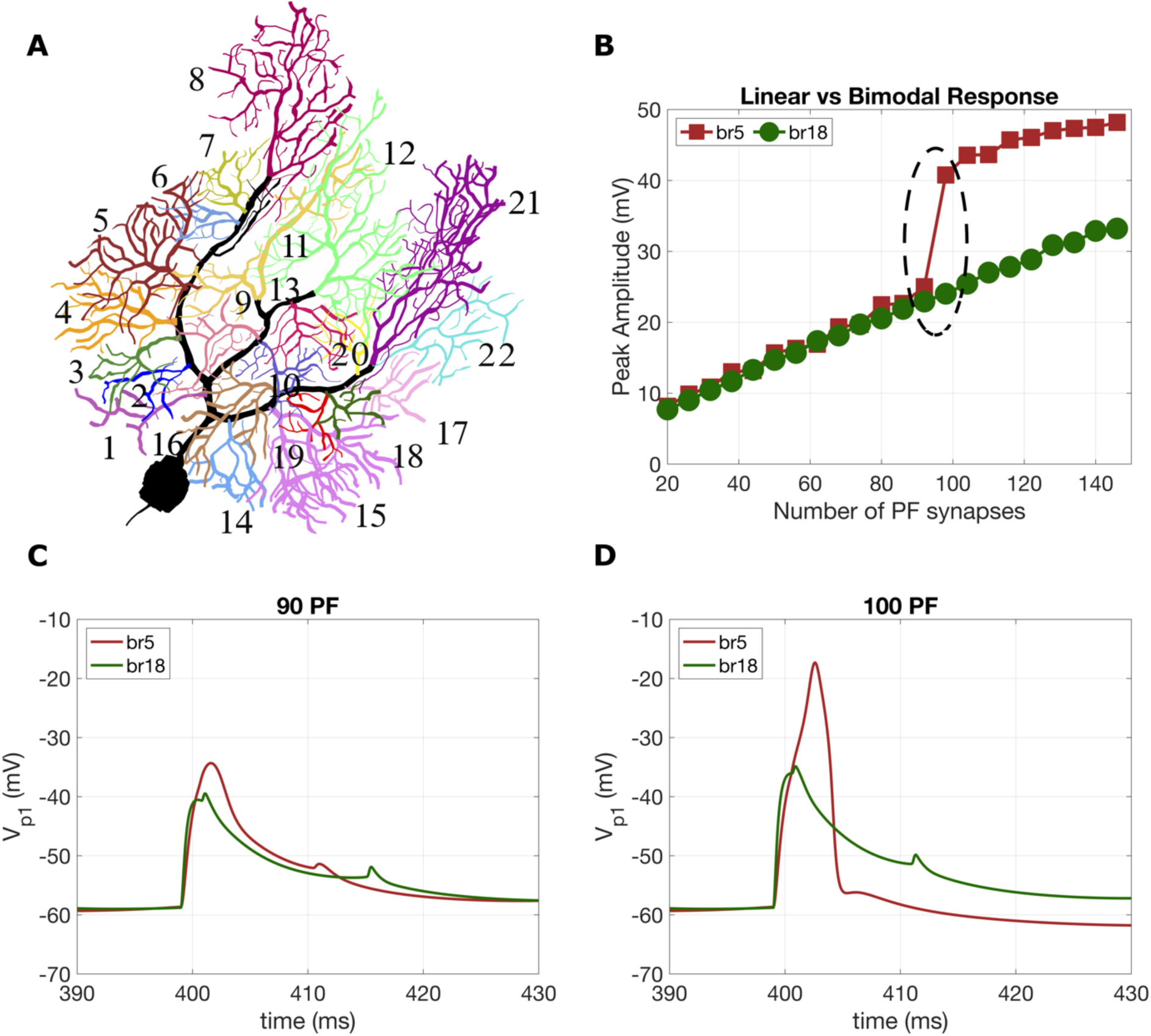
**A.** Branches of the Purkinje cell model. The spiny dendrites were grouped into 22 branches, such that each branch connects to the smooth dendrite shown in the thick black line. **B.** PAR with respect to increasing number of activated PF synapses: bimodal linear-step-plateau response for branch 5 (brown) and linear response for branch 18 (green). Branch 5 is characterized by an initial linear response, followed by a step increase in the gain curve. The “jump” in the peak amplitude is 15mV, occurs for a number of 94 activated PF and is followed by another linear response once the threshold of 96PF is reached. C and D. Voltage responses recorded at the distal point of branch 5 (red) and branch 18 (green), for different PF values: 90PF (panel C) and 100PF (panel D). The response in branch 5 to 100 activated PFs show a dendritic calcium spike is triggered.

### 2.1. Role of Calcium conductance density (CaP) in shifting the gain curve response

We schematically illustrate the linear versus bimodal response in the original model ^27^ in **Figure 1B**, where we use branch 18 to show how dendritic responses linearly increase with increasing activated PF synapses, and branch 5 for showing the bimodal linear-step-plateau response. For branch 5, we observe that initially, dendritic responses linearly increase with increasing activated PF synapses until a large “jump” is triggered (circled in **Figure 1B**). We define as PF threshold, the end point of the jump response, after which the gain curve plateaus. **Figures 1C and D** show the voltage recorded at the most distal point of branch 5 (brown) and branch 18 (green) for 90 and 100PFs, respectively. The corresponding number of PF are simultaneously activated at t=400ms. At the activation time, for 90PFs, we observe a relatively modest increase in both branches: 24.4mV for branch 5 and 18.2mV for branch 18. However, for 100PFs, we observe **in Figure 1D** a large increase of 41.4mV for branch 5, while branch 18 only shows a modest 23.6mV rise. This sharp increase for branch 5 gives the 15mV “jump” shown in **Figure 1B**.

To illustrate how altering the biophysical properties of the PC can result in a change in the synaptic gain curve, we use two different branches with distinct behaviors: branch 3 (see **Figure 2**), for which the PAR increases linearly with PF stimulation, and branch 15 (see **Figure 3**), which already exhibits a bimodal response. For these two branches, we show how the PAR changes when altering the CaP conductance densities (CaP_g) and we analyze how the dendritic spikes propagate.

**Figure 2:**
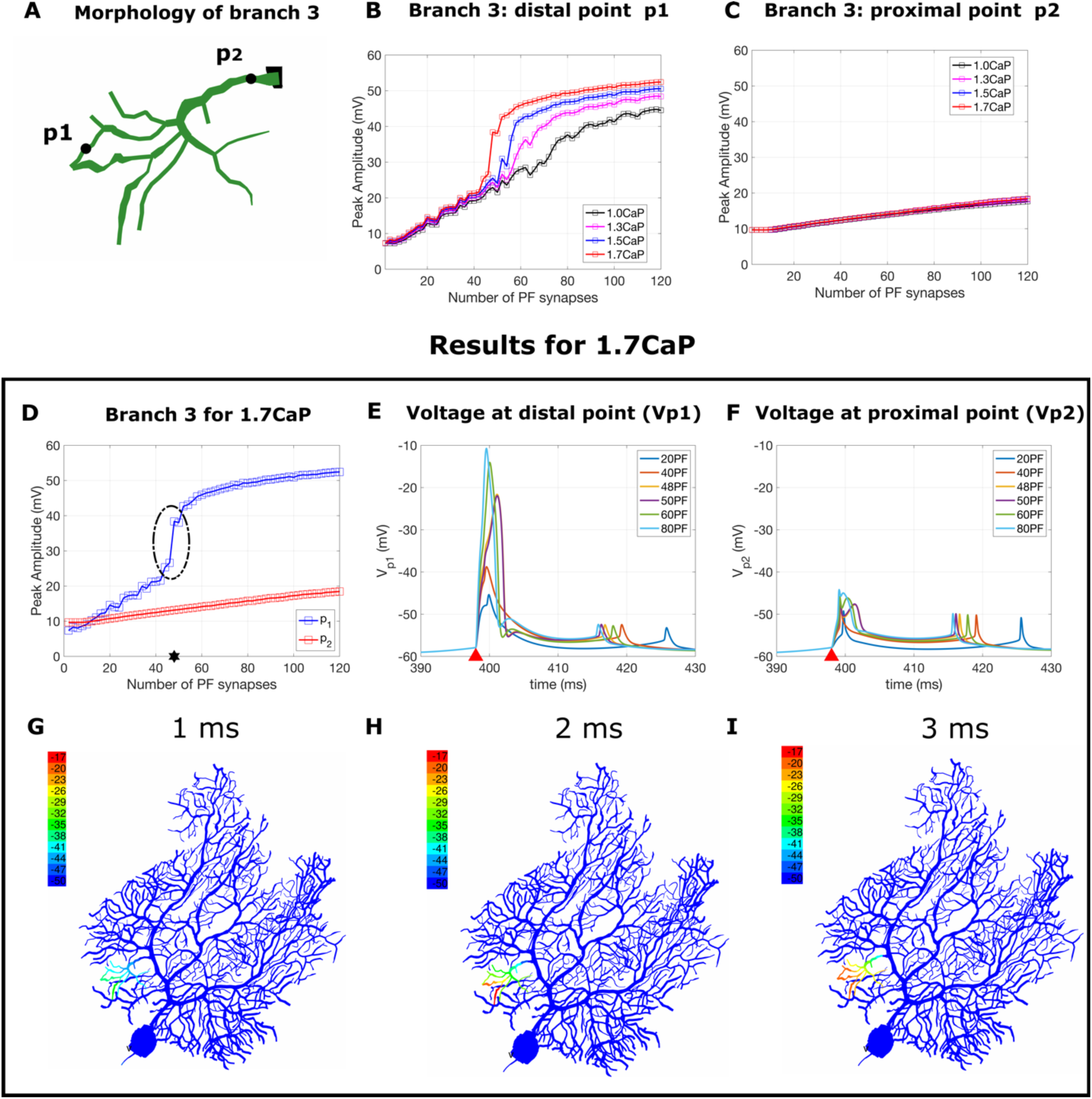
**A.** Illustration of branch 3. The voltage is measured at two points: distal point *p*_1_ and proximal point *p*_2_. **B. and C.** PAR for the distal point *p*_1_ and the proximal point *p*_2_, respectively with increasing number of activated PF synapses. The black line indicates the response when considering the reference value of P-type calcium channel conductance density (CaP_g), while the magenta, blue and red lines show relative increases in P-type calcium channel conductance density of 30%, 50%, respectively 70%. D) PAR for the two points p1 and p2 for an increase of 70% in CaP_g. The bimodal linear-step-plateau response (circled) occurs at a threshold of 48PF (shown using a star). E and F. Voltage response measured at the point *p*_1_ and *p*_2_ for different number of activated PF in [2,120] for the case of 70% increase in CaP_g. The smaller spikelets at the end of each recording are attenuated somatic action potentials. G, H and I. Dendritic spike propagation after 1ms, 2ms and 3ms from activating the PF input. Observe that the dendritic spikes initiate at the tip of branch 3 and propagate towards the smooth dendrite, depolarizing the entire branch. This depolarization is entirely localized, no spreading to neighboring branches is observed.

**Figure 3:**
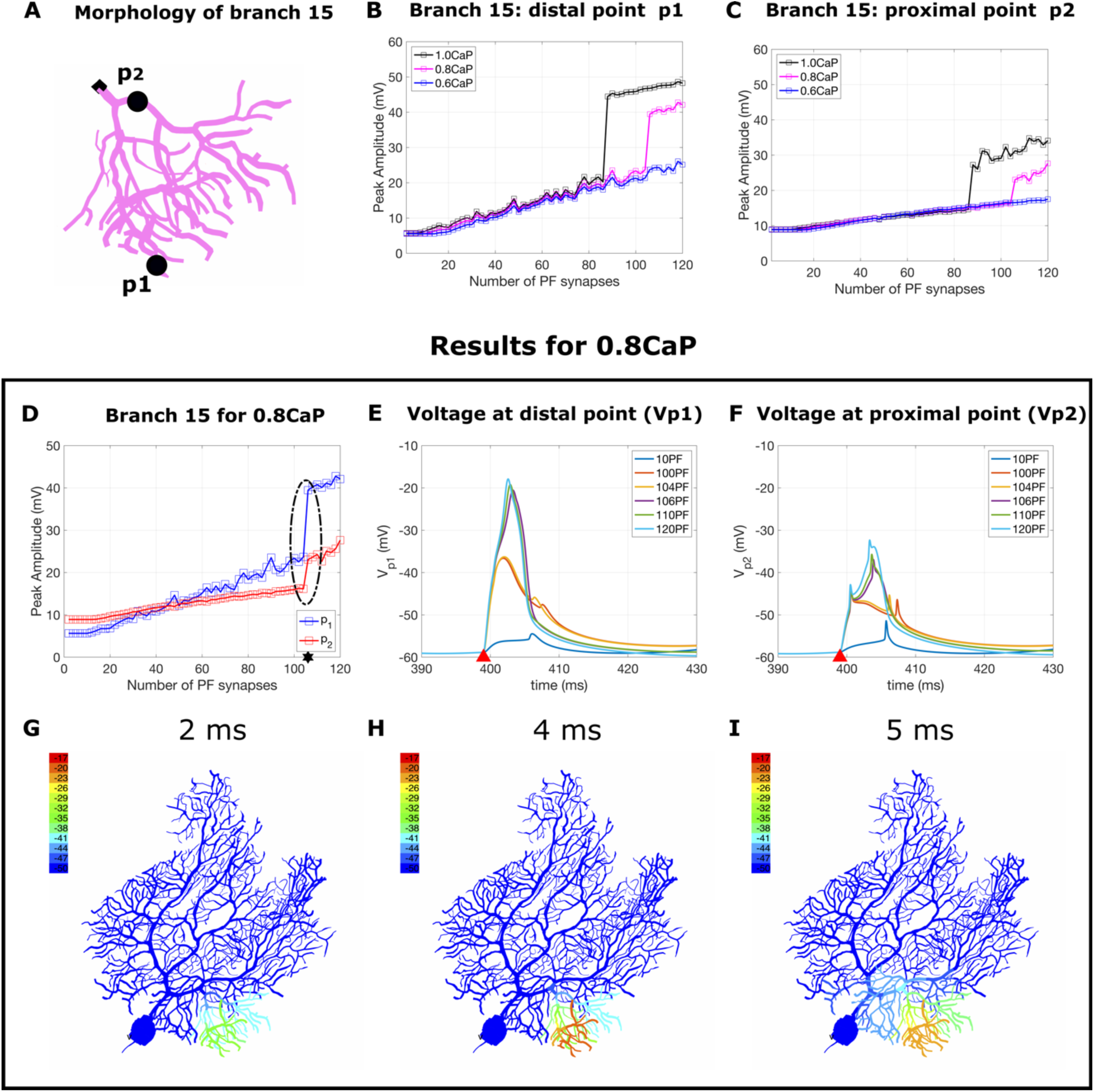
**A**. Illustration of branch 15. The voltage is measured at two points: distal point *p*_1_and proximal point *p*_2_. **B and C.** PAR for the distal point *p*_1_and the proximal point *p*_2_, respectively with increasing number of activated PF synapses. The black line indicates the response for the reference value of CaP_g, while the mangenta and blue lines show relative decreases in CaP_g of 20 and 40 respectively. **D.** PAR for the points *p*_1_ and *p*_2_ for a decrease of 20 in CaP_g. The bimodal linear-step-plateau response (circled) corresponds to a threshold of 106 PF (black star). **E and F.** Voltage response measured at the point *p*_1_ and *p*_2_ for different number of activated PFs in [10,120] for the case of 20 decrease in CaP_g. **G, H and I.** Dendritic spike propagation 2ms, 4ms and 5ms after activating the PF input. Observe that the dendritic spikes initiate at the tip of branch 15 and propagate towards the smooth dendrite, depolarizing the entire branch. In panel I we observe that there is a small depolarization of −41mV of the neighboring branch 14.

**Figure 2A** shows the morphology of branch 3 on which we uniformly distributed 2 to 200 PF synapses with a step of 2PF. We selected two points: a distal point *p*_1_ and a proximal point *p*_2_, in which we recorded the voltage and calculated the PAR with respect to the number of activated PFs. The black line in **Figure 2B**, shows the linear response obtained when using the original uniform ion channel density model.^27^

We gradually increased the CaP_g for the spiny dendrites in branch 3 by 30% (magenta), 50% (blue) and 70% (red). We observed that a 70% increase is sufficient to shift the response from linear to bimodal linear-step-plateau. The “jump” is initiated at 46PF and reaches its plateau at a threshold of 48PF, shown with a black star in **Figure 2D**, and has a peak amplitude difference of 12mV. Unlike the distal point, the PAR at the proximal point *p*_2_ (**Figure 2C**) shows a very small increase when activating the PF input. **Figure 2D-I** considers an increase of 70% in CaP_g.

In **Figure 2E-F**, we visualize the shape of the voltage recorded at the distal point *p*_1_, and at the proximal point *p*_2_ for different number of activated PF synapses. In our simulations, the PC fires spontaneously, as observed experimentally,^37^ and we activate the PF input at time t=398ms. In **Figure 2E**, we observe that activating a sufficiently large number of PF synapses, produces a transition from EPSP to large amplitude dendritic spike when recording the voltage in *p*_1_. On the other hand, the voltage recordings at the proximal point *p*_2_show no significant depolarization (**Figure 2F**).

**Figure 2G-I** show the dendritic spike propagation when activating 48PFs on branch 3. At 1ms after activation, we observe a very small depolarization starting at the tip of branch 3 and propagating towards the smooth dendrite (see Figure 2G). This depolarization increases as the time passes and reaches its maximum depolarization of −17mV after 3ms (see Figure 2I). This depolarization remains entirely local and does not spread to any of the nearby branches. The video of the dendritic spike propagation is available as online Supplementary Material.

When examining a much larger branch such as branch 15 (see Figure 3), we observe a different behavior. This branch already exhibited a bimodal linear-step-plateau response, but it was not reported in the previous analysis^27^ due to its very large threshold at which the “jump” occurs (see black line in Figure 3B), which exceeded the maximum number of PFs considered.^27^ We use this branch as an example of how the bimodal response can in turn be converted to a linear response by reducing the CaP_g. In Figure 3B-C we show the response for the baseline value of 1.0CaP_g used^27^ and we show that even a reduction of 20% of CaP_g conserves the bimodal response (magenta line). However, if we reduce this further by 40%, it shifts to a linear response (blue line).

In panels D-I we show the results obtained for a reduction of 20% in CaP_g. This reduction limits the depolarization onto the neighboring branches, while maintaining the bimodal response circled in Figure 3D, where we observe a very large threshold of 106PF. When examining the PAR in the distal point *p*_1_, we observe a very large “jump” of 15.7mV, while in the proximal point *p*_2_, we have a very modest “jump” of 6.9mV.

The dendritic spike propagation in branch 15 is shown in Figure 3G-I where we activated 106PF. Two ms after activation we observe a small depolarization which starts on the tip of branch 15 and propagates towards the smooth dendrite (see Figure 3G). The depolarization increases further and reaches its maximum after approximately 4ms (see Figure 3H). In Figure 3I we show how, after 5ms from activation, the depolarization minimally spreads to the neighboring branch 14, where it reaches a maximum of −41mV (see video in online Supplementary Material).

### 2.2. Role of Kv4 in blocking the depolarization to nearby branches

The work of Zang and De Schutter^27^ shows that activating clustered PF on branch 8, one of the largest and most distal branches of the PC, produces a very large depolarization onto the nearby much smaller branches 6 and 7. We investigated various modalities of constraining such depolarizations. Our first attempt was to reduce the CaP_g as shown in the previous section. This strategy can indeed decrease the depolarization, however, reducing CaP_g by more than 10% in the case of branch 8 results in a shift of the PAR from bimodal to linear.

Therefore, to limit the depolarization while maintaining a bimodal linear-step-plateau response, we investigated the role of the voltage-activated potassium channel, Kv4. We observed that by increasing Kv4 conductance densities (Kv4_g) on the larger branches and nearby smaller branches, we were able to significantly reduce the spreading. Figure 4D-F shows the dendritic spike propagation for branch 8 when increasing Kv4_g on branch 8, 7 and 6. Notice that in this case, the dendritic spikes propagate throughout branch 8 and they only slightly spread towards branches 6 and 7 (−35mV).

**Figure 4:**
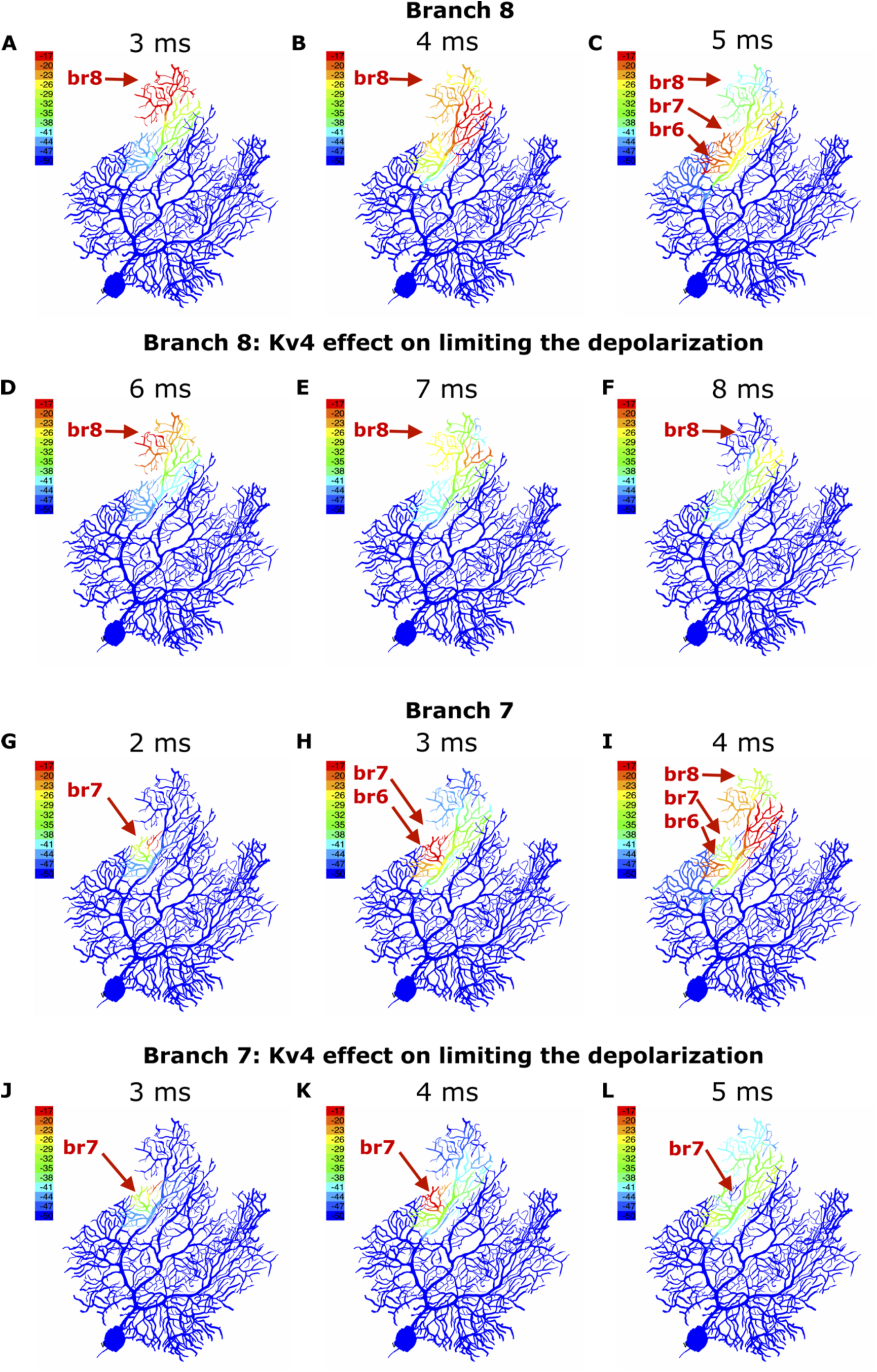
Role of the voltage dependent potassium Kv4 channel in blocking depolarization of nearby branches. **A-C.** Dendritic spike propagation for branch 8 when CaP_g is reduced by 10%. Observe that activating PF on branch 8 produces a very strong depolarization on the smaller neighboring branches 6 and 7. **D-F.** Dendritic spike propagation for branch 8 when increasing Kv4_g by 2.5 fold for branch 5, 2.8 fold for branch 6, 3.5 fold for branch 7 and 3 fold for branch 8. Compared to the previous row (A-C), the spreading onto the nearby branches is significantly reduced. **G-I** Dendritic spike propagation for branch 7. Observe that the dendritic spikes propagate first towards branch 6 and then towards the entire branch 8. **J-L.** Dendritic spike propagation for branch 7 with same changes to Kv4_g as in panels D-F. Notice that the dendritic spikes are entirely constrained to branch 7.

Similar behavior was observed for branch 7 (Figure 4G-I). When activating PF on branch 7, we notice that the dendritic spikes propagate throughout the entire branch (Figure 4G) and fully depolarize the neighboring branch 6 (Figure 4H), after which they spread towards branch 8, provoking a full depolarization (Figure 4I). On the other hand, when increasing Kv4_g (Figure 4J**-L**), we observed that the dendritic spikes initiating in branch 7 (Figure 4J) are entirely constrained within the same branch (Figure 4L). The videos are available in the Supplementary Material.

When examining the entire dendritic tree, we determined that for most branches, the dendritic spikes are entirely constrained within the branch in which they originate. For the branches which exhibited large depolarizations (such as branches 6,7,8,12,13), we adjusted the Kv4_g so that the voltage stayed below −35mV on the nearby branches. All Kv4_g changes are summarized in Figure 5E.

**Figure 5.**
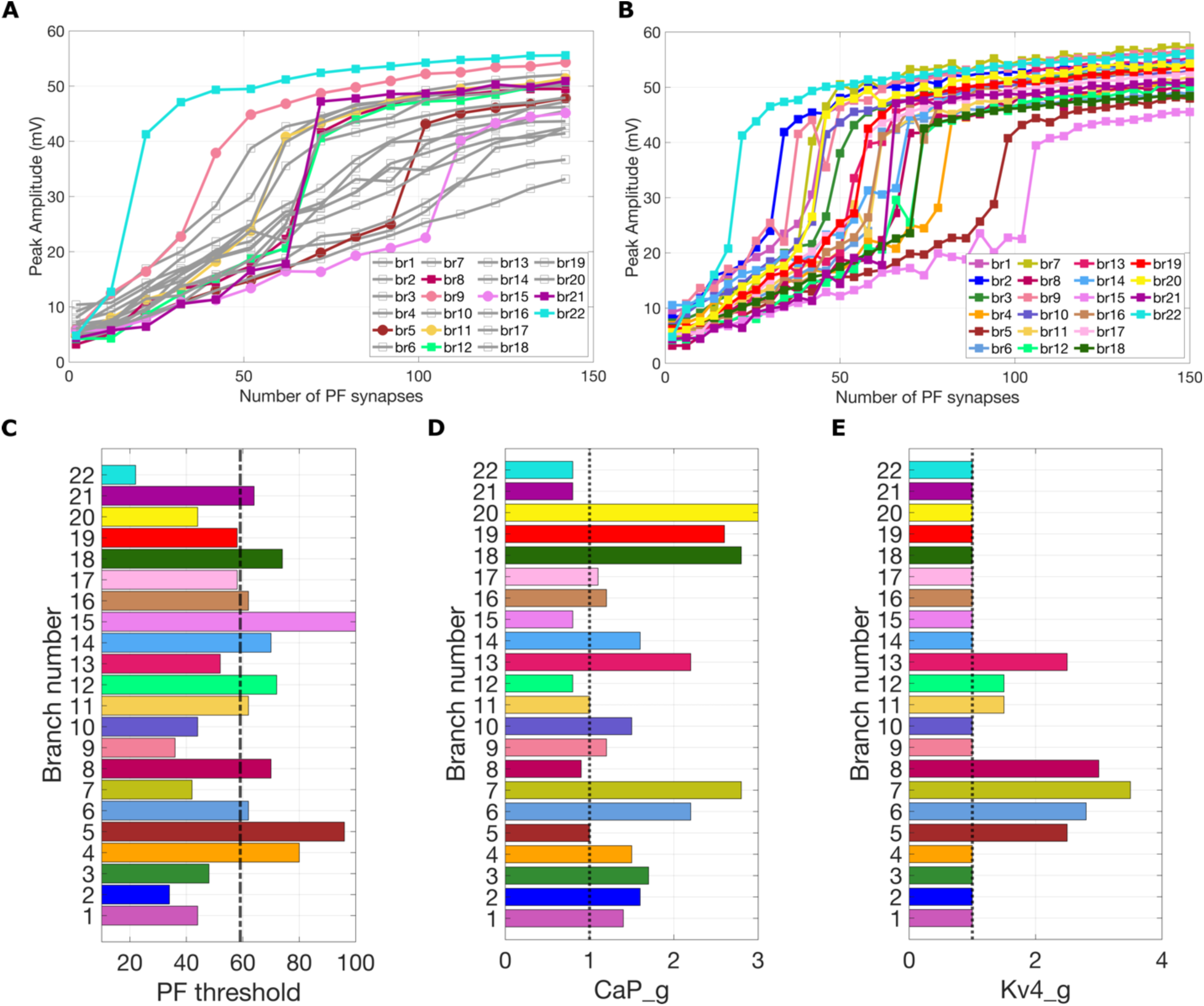
Results and parameters for the heterogenous model: A and B. PAR using the homogenous ion channel model (**A**) or the heterogenous ion density channel model (**B**). The branches shown in gray in panel A exhibit a linear response when simulated using the homogenous model. The branches shown in colors show a bimodal response: branches 8, 12, 21 and 22 (squared markers) were already shown to have a bimodal response^27^ while branches 5, 11 and 15 (circle markers) were overlooked in the previous work due to their very large PF thresholds. Panel B shows that all branches exhibit a bimodal response, when using our heterogeneous ion channel model. **C**. The minimum number of PF required within each branch for initiating the jump. The dashed-dotted line indicates the average threshold. **D and E.** Relative changes in ion channel densities for obtaining the bimodal response for all the branches for CaP_g (panel D) and Kv4_g (panel E). The dotted lines indicate the baseline conductance densities used in the homogeneous model.

We identified two outlying branches whose spreading depolarization could not be fully constrained: branch 12 and branch 18. For branch 12 (see **Figure S6**), we were only partially successful in blocking the depolarization towards the nearby branches 9, 10, 11 and 13. We believe this is due to the special morphology of branch 12, a very large and distal branch located on the center part of the dendritic tree. By adjusting the Kv4_g on branch 11, 12 and 13, we were able to limit the depolarization towards branch 12, when activating branch 11 and branch 13. However, when activating PF on branch 12, the spreading depolarization first reaches branch 13 and 11, after which it continues towards branch 9 and 10. We included the results and discussion regarding branch 12 in the Supplementary material (see **Figure S6**). On the other hand, branch 18 is the smallest branch in our PC (see Figure 6A) and is located in the immediate vicinity of branch 19, being connected to the same smooth dendrite. We believe its morphology, and its close proximity to branch 19 (see **Figure S2**), makes it impossible to contain the depolarization towards its neighboring branches 19 and 17.

**Figure 6.**
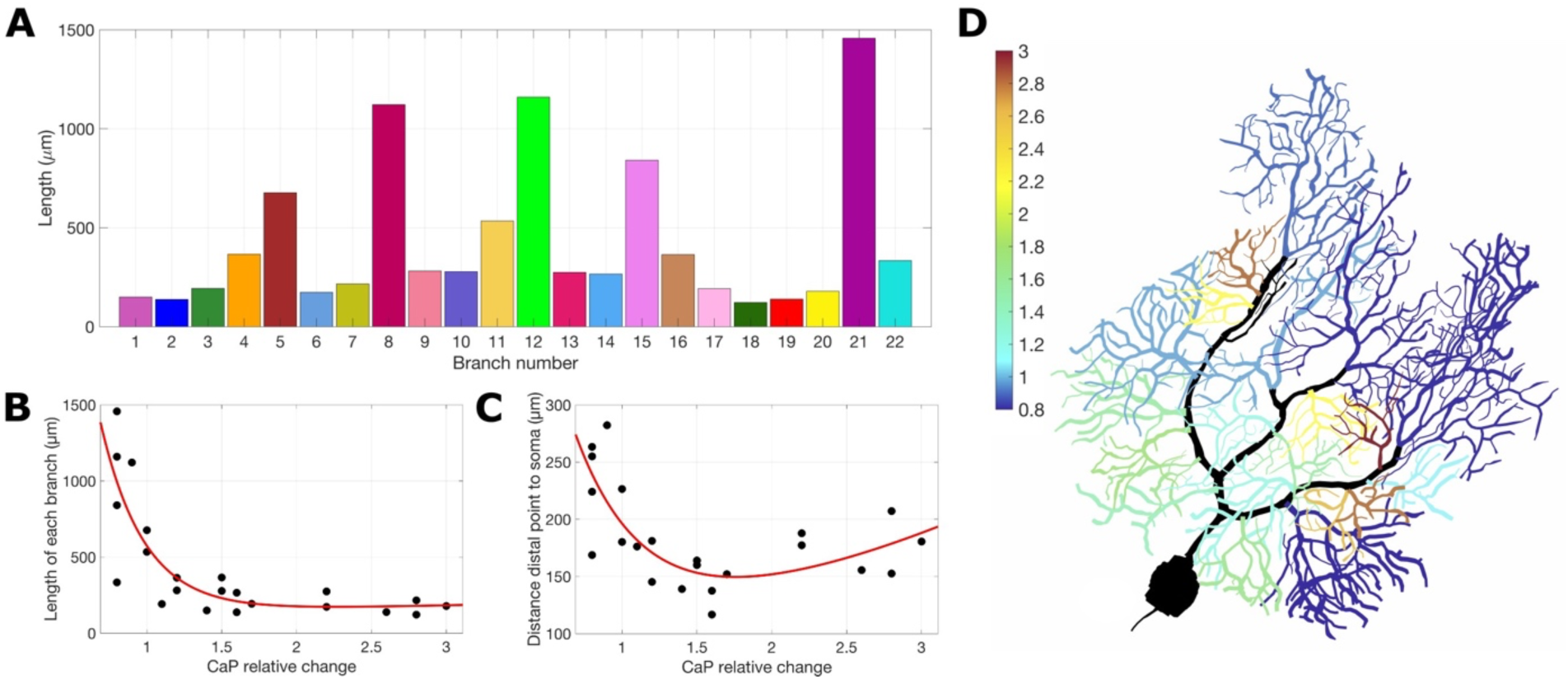
Morphological properties of the branches. **A.** Cumulative lengths of each branch in the PC. **B.** Scatter plots for the relative change in CaP_g and the length of each branch, which were fitted with a sum of exponentials (in red). **C.** Scatter plots of the relative change in CaP_g and the distance from the most distal point of each branch to the soma, fitted with a sum of exponentials (in red). In the first case the R^2^ obtained is 0.6214, while the second one has a R^2^ of 0.5542. **D**. Distribution of relative increase in CaP_g required for obtaining a bimodal response for all branches.

### 2.3. Summary of the results obtained for each branch

Following the same algorithm for each branch in the dendritic tree, we were able to convert the response from linear to bimodal linear-step-plateau for all branches. We summarize the results obtained in Figure 5, where in panel **A** we show the response when considering the original uniform channel density model,^27^ while in panel **B** we show the bimodal response obtained with our novel heterogenous ion channel density model. Panel **A** shows in gray all the branches characterized by a linear response and in color, the eight branches that already showed a bimodal response: in squared markers we indicated the four branches that were captured in previous work,^27^ while in circle markers we show the branches that were overlooked. All our simulations were performed for a maximum of 200PF for all individually activated branches, but as no further jumps were detected after the threshold of 106PF, in Figure 5A-B we show the response till 150PF. The color coding used in all the graphs follows the same color scheme chosen in Figure 1A.

Observe that each branch has a characteristic PF threshold, which we define as the finish point of the “jump” and the starting point of the plateau. We collected these values in Figure 5C, where we observe that the thresholds vary significantly between 22PF for branch 22 to 106PF for branch 15, with an average value of 59PF, expressed by the dash-dotted line in panel **C**. The modifications we made to the CaP and the Kv4 channel densities for each branch are collected in Figure 5D-E and are expressed as relative changes with respect to the baseline values (see **Table S1)**. We correlate the values of the ion channel conductance densities to various morphological properties of the corresponding branch such as length and distance from soma in **Section 2.6**. All the results we show in the rest of this paper follow the parameter choices shown in Figure 5D-E.

In conclusion, in this section, we presented the results obtained when simulating a clustered PF input^2,3^ on each branch of the PC. Our model employing branch-specific conductance densities compensates for the different branch morphologies and achieves a uniform bimodal linear-step-plateau response throughout the PC. In addition, our model fully or partially constrains dendritic calcium spike propagation within the activated branch.

### 2.4. Basic properties of the heterogenous densities model

In response to climbing fiber input, our novel heterogenous PC model generates complex spikes as shown in **Figure S7A**. In addition, we show the frequency-current curve, obtained when injecting different current amplitudes at soma in **Figure S7B.** The results we obtained using the heterogenous ion channel density model (red line in **Figure S7B)** are similar to the F-I curve of the homogenous model^27^ (blue line). Both F-I curves fall in the experimentally measured range (see Figure 1F from Zang et al.^32^).

### 2.5. Quantitative analysis of dendritic calcium spikes

Our simulations capture how strong enough clustered PF input on each branch of the PC, initiate dendritic spikes. Figure 2G-I, Figure 3G-I, and Figure 4 show how the dendritic spikes initiate at the tip of the stimulated branch/branches and propagate towards the smooth dendrite. As they reach the smooth dendrite, their amplitude decreases, as observed experimentally.^2,3^ For most branches in the dendritic tree, the dendritic spikes are either entirely spatially constrained or they provoke minimum depolarization on the nearby branches. The only exceptions are the larger more distal branches such as branch 8 (Figure 4) or branch 12 (**Figure S6**), which strongly depolarize the neighboring much smaller branches 7 and 6 (Figure 4A-C and Figure 4G-I), respectively 11 and 13 (**Figure S6**). This depolarization, which was already captured in the previous work,^27^ was blocked by increasing Kv4 conductance density (see Figure 4D-F and Figure 4J-L) as summarized in Figure 5. Following these modifications, most dendritic spikes remain local within the branch they originate from or, in some instances, provoke minimum depolarization on the nearby branches. However, there are two exceptions (branch 12 and 18) for which our heterogenous model was not fully able to compensate for the distinct morphology of these branches and the propagation of the spikes could be only partially reduced.

We observed that the time required to achieve maximum depolarization is highly dependent on the branch considered, varying from 2-3ms for the small branches situated near the soma such as branch 3 (Figure 2), 10 (**Figure S4**) to 5-8ms for the larger more distal branches such as branch 5 (**Figure S3**), 8 (Figure 4D), 11 **(Figure S5)** or 12 (**Figure S6**).

### 2.6. Morphological factors that influence branch excitability

One possible factor that may control the excitability of each branch is its length. In Figure 6A we show the cumulative lengths of all branches in our dendritic tree. We observe that the more distal branches (8,12 and 21) also have the largest lengths. When correlating the PF thresholds with the length of each branch and the distance from soma, a moderate statistical correlation was found: a positive Pearson correlation coefficient of 0.467 with a p value of 0.028. The correlation between the PF thresholds and the distance to soma was insignificant (Pearson correlation coefficient 0.0483, p value 0.8309).

One other factor that controls the excitability within each branch is the P-type channel conductance density. In order to convert the linear response into a bimodal one, we increased the conductance density of the P-type calcium channels for the spiny dendrites as summarized in Figure 5D. In Figure 6D we show the distribution of CaP conductance densities across the dendritic tree. The color bar indicates the relative increase with respect to the baseline value. As described in the previous sections, the most distal branches such as branch 8, 12, 21 and 22 already exhibited a bimodal response. The CaP_g was lowered to 90% of its value for branch 8 and 80% of its value in case of branches 12, 21 and 22 to limit the dendritic spike propagation from these branches to the nearby branches. Moreover, the relatively large branches such as branch 5 and 11 required no increase in CaP_g. On the other hand, very small branches such as branches 18, 19 and 20 needed a very large increase in CaP_g of 2.8, 2.6, respectively 3-fold of the baseline value.

Following these observations, we analyzed the statistical correlation between the increases in CaP_g required for obtaining bimodal response and two parameters: the cumulative length of each branch (see Figure 6A) and the distance between the distal point at each branch and the PC soma. The length of each branch and the CaP_g relative increase were strongly negatively correlated (Pearson correlation coefficient −0.6246, p value 0.0019). In other words, the larger the branch size, the smaller the relative CaP_g increase needed to obtain the bimodal response. When examining the distance from the distal point of each branch to the soma and the relative CaP_g increase, we obtained a Pearson correlation coefficient of −0.3761 with a p value of 0.084. This moderate negative correlation implies that shorter distances to the soma require larger CaP_g relative increases. Figure 6B-C show scatter plots for the CaP_g relative change with respect to the length and the distance to soma, respectively.

The analysis indicates that the length of each branch is moderately correlated with the PF threshold but highly correlated with the CaP_g increase required for achieving a bimodal linear-step-plateau response.

### 2.7. Multiple branch activation

We simulated co-activation of clustered PF input on two branches, and we analyzed the PAR and the propagation of the dendritic spikes. In Figure 7A-J we show two different examples of co-activated branches, while in Figure 7K-L we describe the results obtained for all possible branch combinations.

**Figure 7.**
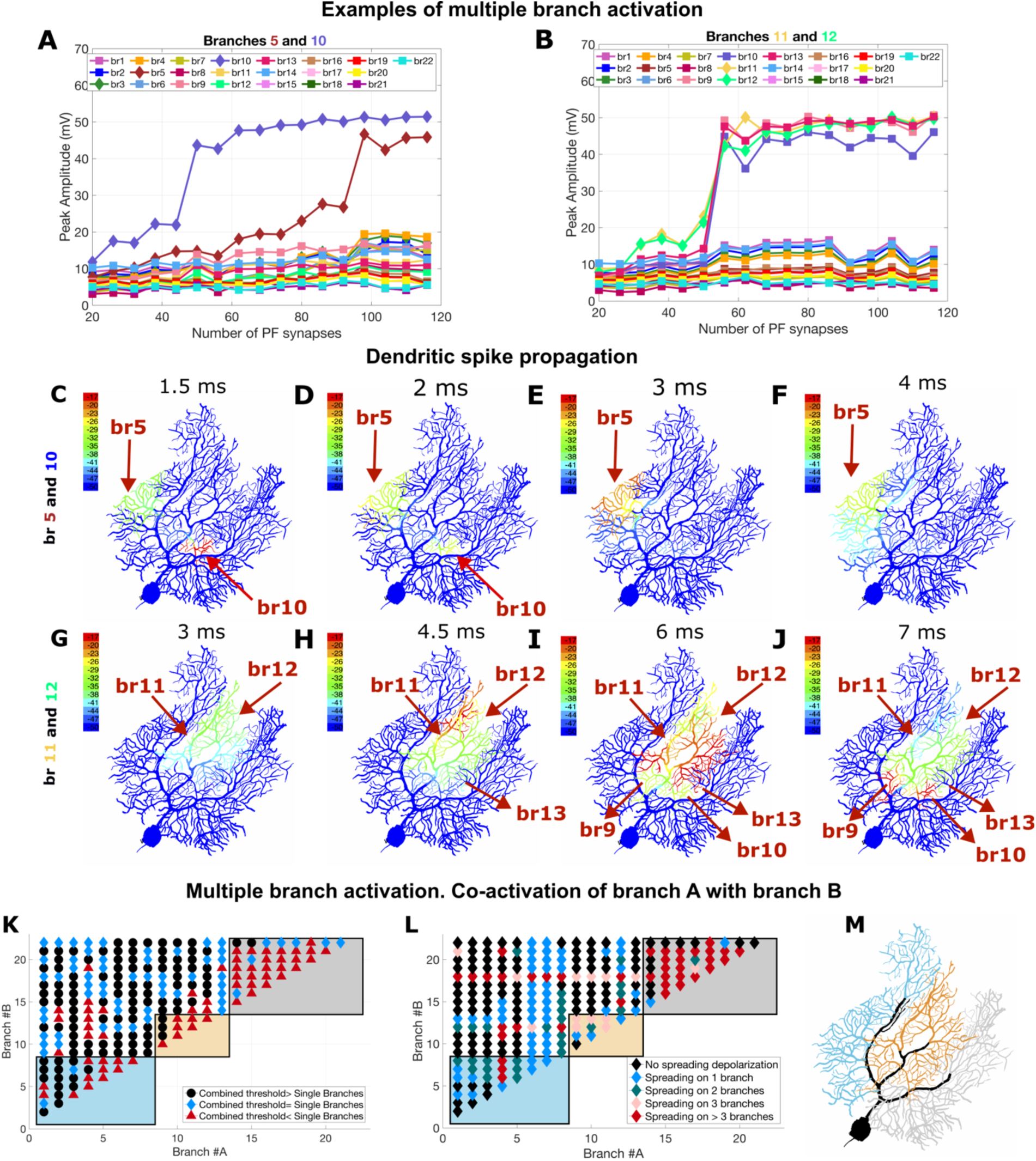
Multiple branch activation: A-B. PAR for simultaneous activation of PF input on branches 5 and 10 (panel **A**) and branches 11 and 12 (panel **B**), respectively. In both cases, we observe that the bimodal response is maintained for the activated branches marked with diamond markers. When co-activating branch 5 and 10, the remainder of the branches exhibit a linear response (squared markers), while when co-activating branches 11 and 12, there is a strong bimodal response in neighboring branches 9, 10 and 13. Panels **C-F** show the dendritic spike propagation for simultaneous activation of branches 5 and 10, while panels **G-J** show the dendritic spike propagation for branches 11 and 12. Observe that for the first example, the dendritic spikes remain localized in the branch they originated from, showing a very small depolarization onto branches 3 and 4 at 4ms (panel **F**) of approximately −41mV. On the other hand, the second case shows a large depolarization (−17mV) on the entire middle part of the dendritic tree (panels **I-J**). **K.** Comparison between PF thresholds for individually activated branches versus co-activated branches. Black markers correspond to at least one of the co-activated branch PF thresholds being larger than that for the individually activated branch, blue markers imply equality while the dark red marker shows that the combined branch threshold is smaller than the PF threshold of the single branch. **L.** Assessment of spreading depolarizations when co-activating different branches. For each pair of co-activating branches, colored markers show how many additional branches have a spreading depolarization that is larger than −35mV. **M.** Main branch divisions of the PC: left division (light blue), middle division (orange) and right division (gray). The colored rectangles in panels K and L indicate that the branch combinations belong to the same main division defined in panel M.

In Figure 7A we activated clustered PF input on two branches: branch 5, located on the left part the dendritic tree and branch 10, located on the middle part of the tree. We uniformly distributed 20 to 120PF on each of the two branches. In Figure 7A we observe that the bimodal response is maintained for the two activated branches, while the remainder of the branches show no increase in their peak amplitude. When simultaneously activated, branch 5 reaches its step response for a threshold of 94PF, while branch 10 only requires 46PF. These PF thresholds are very similar to those obtained when individually activating each branch: 96PF for branch 5 and 44PF for branch 10 (see **Figure S3** and **S4**).

The local activation of the two branches can be better seen in Figure 7C-F, where we visualize the dendritic spikes which initiate at the tip of each branch, propagate within these branches, and achieve maximum depolarization in Figure 7C for branch 10, after only 1.5ms from activation and in Figure 7E for branch 5, which requires approximately 3ms. In Figure 7F we see a very small depolarization produced by branch 5 onto its neighboring branches 4 and 3, while the dendritic spikes originating in branch 10 remain entirely constrained.

**Figure 7B** shows the co-activation of branches 11 and 12, two neighboring branches located on the distal middle section of the PC (see Figure 1A). Unlike the previous branch combination shown in Figure 7A, we see that once the threshold of 54PF is reached, a large “jump” occurs not only for branches 11 and 12 but also their neighboring branches 9, 10 and 13. This threshold is significantly smaller than the scenario in which the two branches are individually activated: 62PF for branch 11 (**Figure S5**) and 72PF for branch 12 (**Figure S6**).

**Figure 7G-J** show the dendritic spike propagation when co-activating branch 11 and 12. We observe how initially branch 11 and 12 are activated, after which the depolarization spreads on the lower branches 9, 11 and 13, where it reaches −17mV. The co-activation of branches 11 and 12 is one of the most extreme scenarios in our analysis, in which dendritic spikes spread on the entire middle main branch. We believe this occurs due to the very close proximity of the two branches (see **Figure S2**), their large sizes (see Figure 6A), and their more distal positions.

**Figure 7A** and **B** show two opposite examples of multiple branch co-activation. In order to examine the responses in the entire dendritic tree, we analyzed all possible two by two branch co-activations and we performed two sets of tests (Figure 7K-L). To ease the description of the results, we show in Figure 7M three main divisions of the PC: the left most main division (in light blue), the middle main division (in orange) and the right main division (in gray). The squared rectangles colored in light blue, orange and gray, respectively in Figure 7K-L indicate co-activated branches that belong to the same main division.

First, we compared the threshold for co-activation of the two branches with the thresholds of individually activated branches. We observed that for most of the tree, the co-activated branch thresholds are larger or equal than the threshold of the single branches (black and blue markers in Figure 7K). For example, the co-activation of branches 5 and 10 (see Figure 7A) corresponds to this case. However, when activating nearby branches or branches belonging to the same main division (indicated in the colored rectangles in Figure 7K), due to the spreading depolarizations between the two branches and to the surrounding branches, the combined threshold becomes smaller than the individually activated branch threshold (dark red markers in Figure 7K). The extreme scenario of branch 11 and 12 described above corresponds to this second case. In conclusion, the branches act as individual processing units if they are well separated from each other, ideally located on different main divisions. The closer the co-activated branches are to each other, the smaller their PF threshold becomes, and they no longer function as individual units.

Second, for all possible branch combinations, we studied how their dendritic spikes propagate. In Figure 7L we show whether their dendritic spikes remain local, or they spread towards the neighboring branches. In case of spreading depolarization, we counted how many neighboring branches show a voltage larger than a set threshold value of −35mV. The black diamond symbol in Figure 7L shows no spreading depolarization, while the dark red marker shows that the depolarization is detected in more than 3 branches. While most of the dendritic tree shows no significant depolarization, there are few exceptions that stand out. Firstly, branch 6 co-activated with any other branch (blue diamond marker), results in a depolarization of its neighboring branch 7. Conversely, branch 7 triggers a depolarization on branch 6. This suggests that these two very small branches that are extremely close to each other (see **Figure S2**) are intertwined and could be redefined as a single branch. The same occurs with branch 18 and 19, the smallest branches of the PC (see Figure 6A), which are also very close. As we discussed in **Section 2.2**, we were unsuccessful in containing the spreading depolarization when individually activating branch 18, as it contacted its close neighbor branch 19 and further spread on the right main division. Here, when co-activating branch 18 with any of the branches in the PC, we observed more extreme depolarizations. Also, significant depolarizations occur when co-activating branches from the center main division (orange rectangle), because, as soon as branch 12 is co-activated with any other branch, there is significant depolarization on its neighboring branch 11 and 13. This was also discussed in **Section 2.2** and in the Supplementary **Figure S6**, in which we show how the dendritic spikes propagate when activating PF only on branch 12. Of course, the dendritic calcium spikes originating in this branch, when co-activated with any other branch, trigger stronger depolarizations.

Note that for this analysis, we maintained the same Kv4_g parameter choice as defined in Figure 5E and **Table S2**. Further increasing Kv4_g for the co-activated branches could possibly have a significant effect in reducing these depolarizations. However, our interest here was to determine whether these branches can function as individual units, based on morphological criteria.

## 3. Discussion

### 3.1. Building the first heterogenous ion channel density model

In vitro studies have shown that dendritic calcium spikes can be triggered by clustered PF input.^2,3^ These dendritic spikes are caused by the activation of dendritic voltage-gated calcium channels. Confirmation of their study in vivo has been very challenging using conventional patch-clamp recordings, but recently, with the development of techniques such as in vivo calcium imaging, various authors recorded PF evoked dendritic spikes.^38–42^ In particular, Roome and Kuhn^42^ have recently proposed a fast two-photon imaging technique that simultaneously records voltage and calcium signals from the spiny dendrites of PCs in awake mice and described dendritic spikes triggered by strong PF input.

Numerous recent studies, both experimental^2,33,35,36,43^ and computational,^27,32^ have brought significant evidence supporting a heterogenous dendritic excitability of PCs. The underlying mechanisms which determine the inhomogeneous excitation across the different dendritic branches are associated with various ion channels such as SK^33,35,36^, BK^2^, or A-type K channels.^32,44^

So far, the classical simplifying approach used in PC computational models was to assume that all ion channel densities are uniform in the spiny dendrite.^8–10,27,29–32,45–47^ There are, however, a few pyramidal cell models, in which dendritic ionic channel densities are defined with respect to the path distance from soma.^48,49^

In light of recent evidence highlighting heterogenous dendritic excitability,^2,33,35,36,43^ we propose, to our knowledge, the first heterogenous ion channel density PC model, in which each dendritic branch is characterized by its own set of ion channel conductance densities. Our work is based on two previous models,^27,32^ and continues on the pioneering PC model of De Schutter and Bower^8–10^. For a deeper understanding of how PC models were created, how they evolved over the last fifty years and how they were able to predict significant dendritic properties, we direct the reader to a review article^11^. For a broader understanding, not limited to the cerebellum, of how computational models contributed to uncovering dendritic functions, we recommend a recent review which highlights the importance of developing intertwined computational and experimental methods.^50^

Our heterogenous PC model, as many other models,^27,32^ assumes a uniform PF distribution, with synapses distributed uniformly across each activated branch. This common simplifying modeling approach will be readdressed in future work due to evidence that more distal branches might possess a significantly higher spine density^51^ with each dendritic spine corresponding to one PF input. The proximal dendrites are thought to have less spines due to competition with climbing fibers, which innervate the proximal dendrite.^52^

### 3.2. Heterogenous parallel fiber thresholds across the dendritic tree

In agreement with previous work^27^, our model predicts that in the absence of a sufficiently strong clustered PF input, dendritic responses have a linear increase with PF synapses for all branches.^53^ When a specific threshold is met, the dendritic responses exhibit a jump caused by a dendritic calcium spike, followed by a plateau. Using our heterogenous model, we obtained such a bimodal response throughout the entire dendritic tree. However, the PF thresholds, which represent the minimum number of PF input required to produce a dendritic calcium spike, significantly differ depending on the branch (see Figure 5C). The large difference between the thresholds denotes a strongly inhomogeneous excitability due to specific morphology of the dendritic tree which requires further study. Interestingly, similar PF thresholds apply to multiple branch activation when selecting co-activating branches (Figure 7K) such that they belong to different main divisions (Figure 7M). This indicates that although the branches were defined based on morphology, they can act as independent computational units. However, the same does not hold when co-activating neighboring branches or branches located within the same main division where we observed that the PF threshold is significantly lower than when individually activating one branch.

## 4. Conclusions

Our pioneering heterogenous ion channel density model is the first proposed, to our knowledge, not only in the cerebellum but in the whole brain. Each branch of our PC is characterized by its own set of ion channel conductance densities. Dendritic branch-specific generation of calcium spikes has been intensively studied in the recent years due to its important role in synaptic plasticity and in facilitating information storage.^21,24,54,55^ Our simulations show how altering the biophysical properties leads to a change in the synaptic gain curve from linear to bimodal linear-step-plateau and vice-versa, suggesting that each branch can multiplex at a cellular level. Modulation of intrinsic excitability at a dendritic branch level was proposed in many recent articles via different ion channels such as SK^33,35^ or A-type potassium channels.^32,44^ The immense impact that the intrinsic excitability plasticity of dendritic branches plays in cerebellar learning has been discussed in detail in the work by Ohtsuki et al.^34,36^ Our study is the first to suggest that P-type Calcium channel can modulate PAR at single branch level. CaP channels are of utmost importance in the cerebellum, they are widely distributed on PC somata and dendrites^56,57^ and mutations in their alpha_1A protein-pore forming subunit have been linked to various neuropathologies^58–60^ such as ataxia, migraine or epilepsy.

Using our heterogenous model, by tuning CaP conductance densities for each branch, we obtained a uniform response throughout the cell. Our results show how sufficiently strong activated PF input on each branch produces a dendritic calcium spike. However, these PF thresholds vary considerably between the different branches and therefore suggest a strong inhomogeneous excitability across the tree. We examined how morphological properties can affect the excitability of each branch and we discovered that the length of each branch is strongly correlated with the increase in CaP required for triggering the bimodal response. This explains why the larger, more distal branches, do not require any additional increase in P-type calcium channel conductance density and already exhibited a bimodal response.^27^

By modulating ionic channels at each branch, we were able to compensate for the different branch morphologies, obtaining a uniform response throughout the PC. When studying dendritic spike propagation, we observed that for most branches, the depolarization remains localized in the branch in which PF were activated. The much larger and distal branches, however, tend to depolarize on the nearby significantly smaller branches. For these cases, we observed that increasing Kv4_g blocks the spreading of the dendritic spikes. The role of Kv4 channel in promoting or suppressing the spread of dendritic calcium spikes was previously discussed^32^ for the case of complex spikes.

When co-activating different branches of the PC, we observed that most branches function as individual processing units, having a PF threshold equal or larger than the individually activated branches and the propagation of their dendritic calcium spikes is relatively constrained. This leads us to speculate that PCs may be able to actively control and enhance their capacity of information processing.

## 5. Materials and Methods

Our model is a modification of a PC model proposed by Zang and De Schutter,^27^ which consists of four parts: the soma, axon initial segment, smooth dendrites, and spiny dendrites. The initial conductance densities for the four compartments are given in **Table S1**. All simulations were implemented in NEURON 8.2, while for the analysis and plots we used MATLAB R2022a. The codes are available on ModelDB.

The model encompasses many different ion channels such as voltage-dependent sodium channels, voltage-dependent potassium channels (Kv1, Kv4, Kv4s), voltage dependent T-type and P-type calcium channels (CaP), large conductance calcium activated potassium channels (BK), small conductance calcium activated potassium channels (SK2). These channels were added to the model in agreement with recent experimental findings and were discussed in detail in the previous work by Zang et al.^27,32^ In this work, we focus on the voltage dependent P-type calcium channel whose maximum conductance we alter for each dendritic branch of the PC, in order to obtain a bimodal response. In addition to the voltage dependent CaP_g, we have also modified Kv4_g for blocking the depolarization of the more distal branches onto the smaller neighboring branches. All conductance densities used in the heterogenous model are shown in Figure 5 panels D and E and in **Table S2**.

As a first step in our analysis, we extended the number of activated PFs distributed for each branch from a maximum of 80 to 200. The activated PFs are distributed uniformly with respect to the length of each branch.^27^ This extension led us to observe that, in addition to the four branches whose bimodal response was previously observed,^27^ there are more branches (5, 9, 13 and 15) that exhibit similar response (see Figure 5A).

### Parameter selection goals

In developing the first heterogeneous ion channel density model, our main goal was to achieve a bimodal linear-step-plateau response for each activated branch, while constraining the dendritic calcium spikes propagation. Our parameter selection was done such that:

1. Each activated branch shows a bimodal response with a minimum −10mV jump.
2. The dendritic spikes originating in each activated branch remain relatively constrained within the branch and do not propagate towards the neighboring branches (V>-35mV)

For each branch in the dendritic tree, we selected minimum two points: one distal point and one point located in the vicinity of the smooth dendrite. We calculated the PAR with respect to increasing numbers of activated PFs and we determined whether they show bimodal linear-step-plateau or linear response. For the branches which exhibited a linear response, we gradually increased the CaP_g until a bimodal response was obtained. All the other parameters were kept constant as defined in **Table S1**. We selected the minimum CaP_g value increase required to obtain a bimodal response with a jump of minimum −10mV. For each of these simulations, we analyzed how the dendritic spikes propagate for different number of activated PFs and whether this response remains local or propagates to the nearby branches. In the cases where the propagation onto the nearby branches was very large, we blocked it using increases in Kv4 maximum conductance.

The largest PF threshold at which any branch attains its bimodal response is 106PF and corresponds to branch 15. All simulations were run for PF numbers between 10 and 200, with a step of 2PF. As no jumps were detected after 106PF, when plotting the results, we used a maximum of 120PF for simplicity.

When analyzing the morphological properties of the branches (Figure 6B and C) we used Matlab curve fitting toolbox to find the best fit for our data: a sum of exponentials *f(x) = a ⋅exp (bx) + c ⋅ exp (dx)*, with coefficients *a* = 1.34*e* + 04, *b* = −3.444, *c* = 130 *and d* = 0.01139 for panel **B** and *a* = 801.5, *b* = −2.207, *c* = 82.18 and *d* = 0.2742 for panel **C**.

Due to the small step of 2PF considered between simulations, and the uniform distribution of PF synapses for each number used, we sometimes observed small noise in the PAR with increasing PF synapses (see for example **Figure S5B**). This was not captured before due to the much larger step used.^27^ However, the small step allowed us to determine more precisely the PF threshold and compare it to the threshold when co-activating two branches. The branch co-activation was done by simultaneously activating the same number of PF synapses on two different branches at t = 400ms. Each pair of co-activated branches have same parameters (Kv4_g and CaP_g) as in the individually activated scenario (given in Figure 5). When analyzing the spreading depolarizations in the case of co-activated branches (Figure 7L), we counted how many additional branches, other than the 2 activated branches, show depolarizations larger than a set threshold of −35mV. Due to the noise described above, we imposed the additional condition that this condition must be met for activation of minimum 5 different numbers of PF.

## Acknowledgements

This work is supported by the Okinawa Institute of Science and Technology Graduate University.

## Author Contributions

GC and EDS conceived this study. GC performed all simulations. GC and EDS analyzed the data. GC and EDS wrote the manuscript.

## Declaration of interests

The authors declare no competing interests.

## Supplementary Material

### 1. Simulation of clustered PF input on branch 15 and branch 8 for the reference value of CaP_g

The first section of the supplementary material aims to complement the results shown in Figure 3, where we discussed branch 15 for a 20% reduction in CaP_g. Such reduction was done to limit the depolarization to the nearby branches. **Figure S1**, shows the same branch 15, this time with the baseline value of CaP_g. **Figure S1A** displays a stronger “jump” of 24.2mV, occurring at a lower PF threshold of 88PF while **Figure S1D-F** show the dendritic spike propagation. Observe that in this case, panel F captures a much stronger depolarization on the neighboring branch 16, where the depolarization reaches −35mV, and branches 18 and 19 where it reaches −41mV. Moreover, we can observe the dendritic spikes travelling even further up the dendritic tree, causing very small depolarizations of −44mV on branches 17 and 20.

In the Results section we used branch 8 to show how the propagation of dendritic spikes can be suppressed by increasing Kv4 conductance density (Figure 4). Here, we show the detailed results for this branch. **Figure S1G-L** shows the morphology of branch 8 and the two selected points: a distal point *p*_1_ and a proximal point *p*_2_. Branch 8 is part of the four branches that already exhibited a bimodal linear-step-plateau response,^27^ as it can be observed in panel B in black line. Observe that reducing the CaP_g by 30% (red line), converts this response into a linear one. Reducing the CaP_g by 10% (magenta line), maintains its bimodal response. In panels J-L we show the results for a 10% CaP_g reduction, which lead to a jump of 16.5mV and a PF threshold of 70.

**Figure S1:**
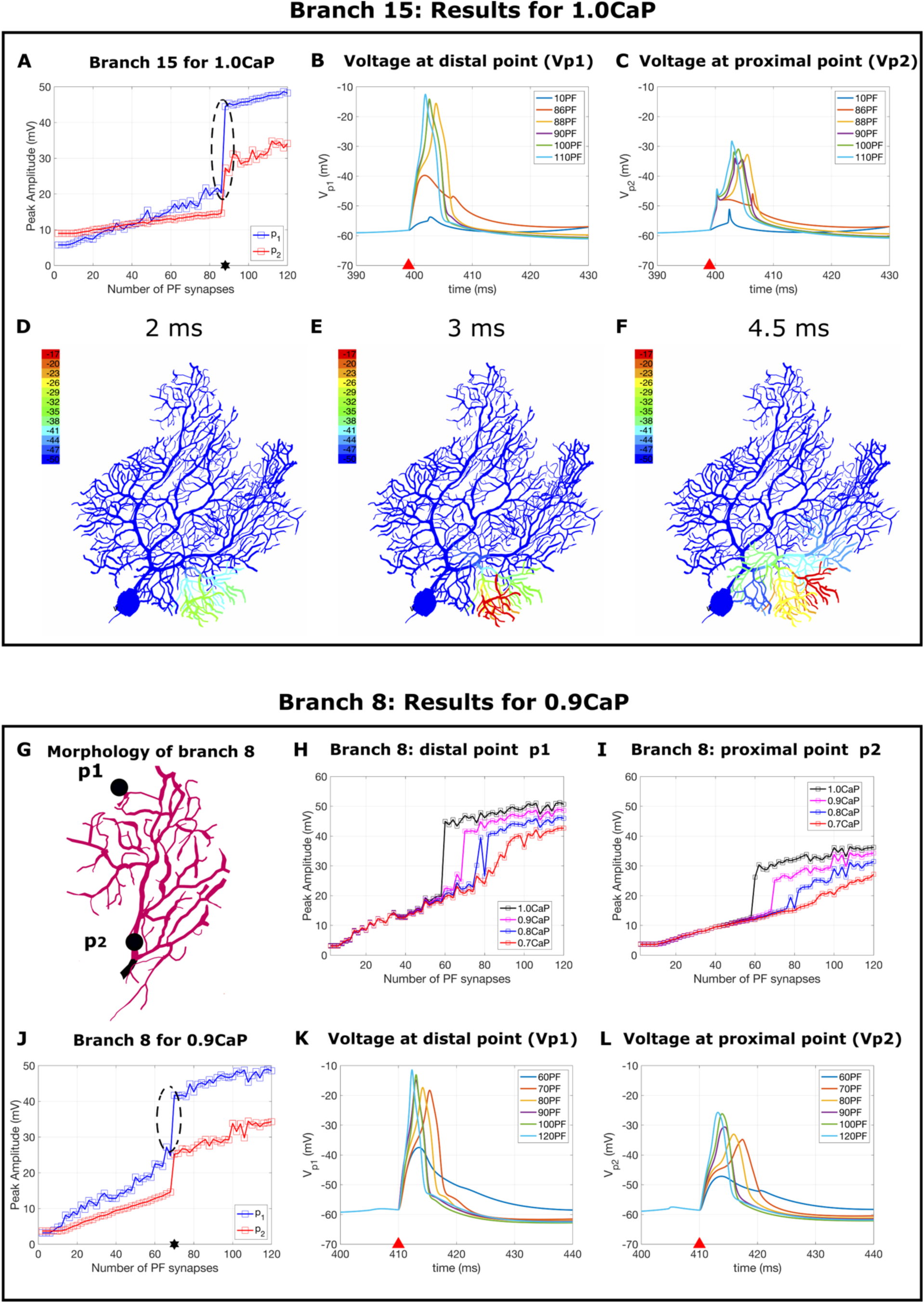
Branch 15 for the reference value of 1.0CaP_g and Branch 8 for 0.9CaP. A. PAR for the distal point *p*_1_ and for the proximal point *p*_2_. The bimodal linear-step-plateau response (circled) occurs at a threshold of 88 PF (shown using a star). B and C. Voltage response measured at the points *p*_1_ and *p*_2_ for different number of activated PF in [10,120]. D, E and F. Dendritic spike propagation 2ms, 3ms and 4.5ms after activating the 88PF. Observe that the dendritic spikes initiate at the tip of branch 15 and propagate towards the smooth dendrite, depolarizing the entire branch. Unlike Figure 3, where we reduced CaP_g by 20%, observe that in this case, the dendritic spikes depolarize nearby branch 16 up to −35mV, produce depolarization of −41mV on branches 18 and 19 and also slightly depolarize branches 17 and 20. G. Illustration of branch 8. The voltage is measured at two points: a distal point *p*_1_ and a proximal point *p*_2_. H and I. PAR for the distal point *p*_1_ and for the proximal point *p*_2_, respectively with increasing number of activated PF synapses. The black line indicates the response when considering the reference value of P-type calcium channel conductance density (CaP_g), while the magenta, blue and red lines show relative decreases in P-type calcium channel conductance density of 10%, 20%, and 30% respectively. J. PAR for the two points for a reduction of 10% in CaP_g. The bimodal linear-step-plateau response (circled) occurs at a threshold of 70 PF (shown using a star). K and L. Voltage response measured at the points *p*_1_ and *p*_2_ for different number of activated PF in [10,120].

### 2. Distances between branches

In Figure 4, we showed how dendritic spikes propagate and how increasing Kv4 can be used to block their spreading. In Section 2.2 we discussed the few branches for which, when individually activated, we obtain large spreading depolarizations such as branch 12 or 18 and we theoretized that this might be due to very small distances between the activated branch and its neighboring branches.

**Figure S2** complements this discussion and shows the distances across the smooth dendrite between all the branches in the tree. As expected, we observe that one of the smallest distances between neighboring branches is between branch 18 and 19 (2.03 µm, dark red marker), which explains the very strong spreading depolarization we discussed. The other dark red marker in **Figure S2** corresponds to the distance between branches 9 and 10, which although very small, does not result in any spreading depolarization when PF clustered input is activated on either of the two branches. This is due to the fact that these branches are in fact located on opposite sides of the smooth dendrite. The next smallest distance in the tree corresponds to branches 6 and 7 (6.67 µ m, pink marker), which we also addressed in Section 2.2, and whose spreading depolarization we managed to contain by increasing Kv4.

**Figure S2:**
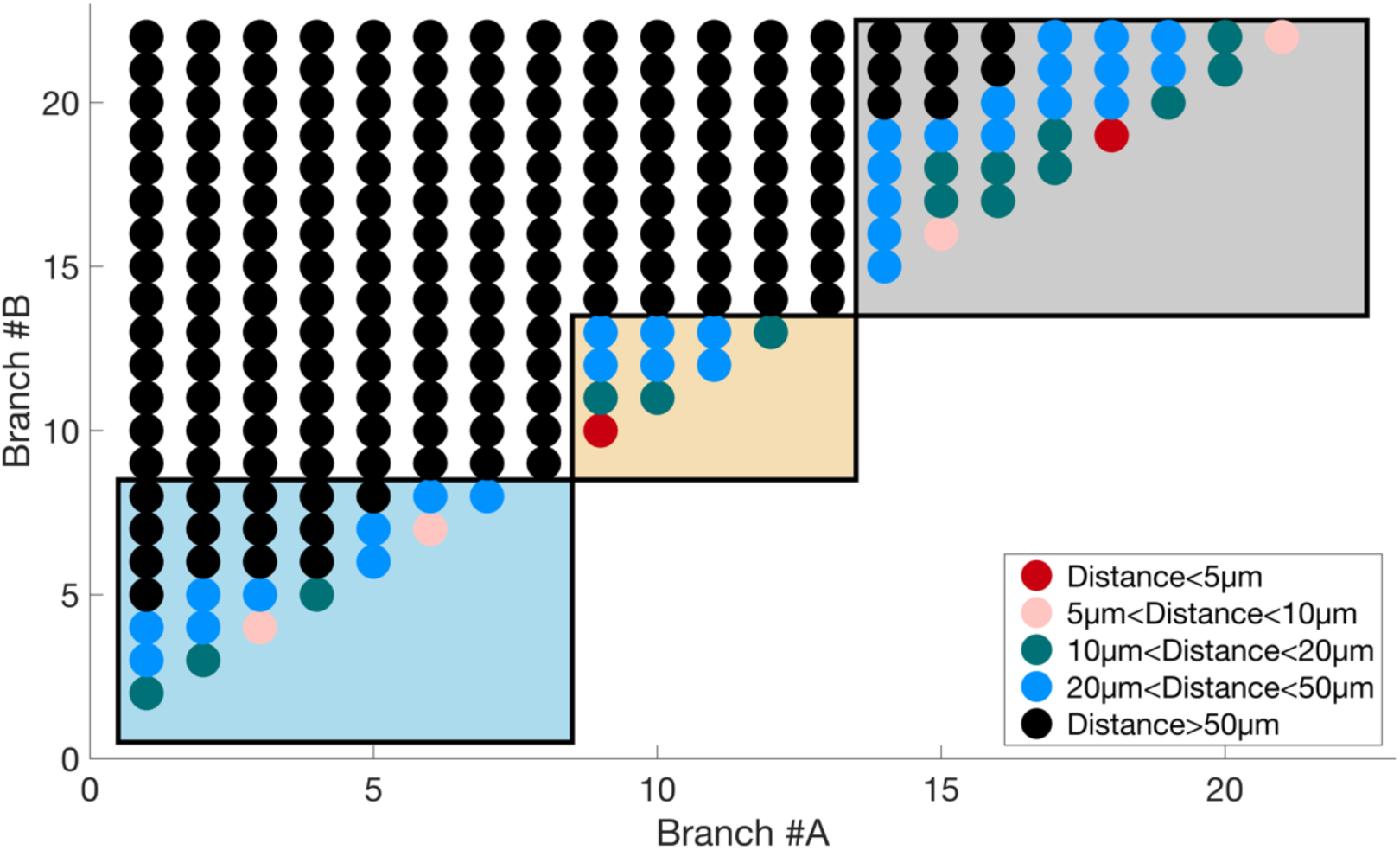
Distances between branches. The dark red marker indicates that the distance between branch A and Branch B is smaller than 5 *µ*m, while the black markers show distances larger than 50 *µ*m.

### 3. Individual clustered PF activation on branches 5, 10, 11 and 12

The following figures aim to complement Figure 7, where we presented two examples of co-activated branches: branch 5 and 10 (Figure 7A) and branch 11 and 21 (Figure 7B). **Figures S3-S6** display the results when individually activating clustered PF input on these branches: branch 5 (**Figure S3**), branch 10 (**Figure S4**), branch 11 (**Figure S5**) and branch 12 (**Figure S6**). For each branch mentioned above, we show the points at which we recorded the membrane potential, the PAR at the distal point and proximal point for different values of CaP_g and the recordings of the voltage recorded at soma, at the distal and proximal points. Furthermore, we present how the dendritic calcium spikes propagate.

**Figure S3:**
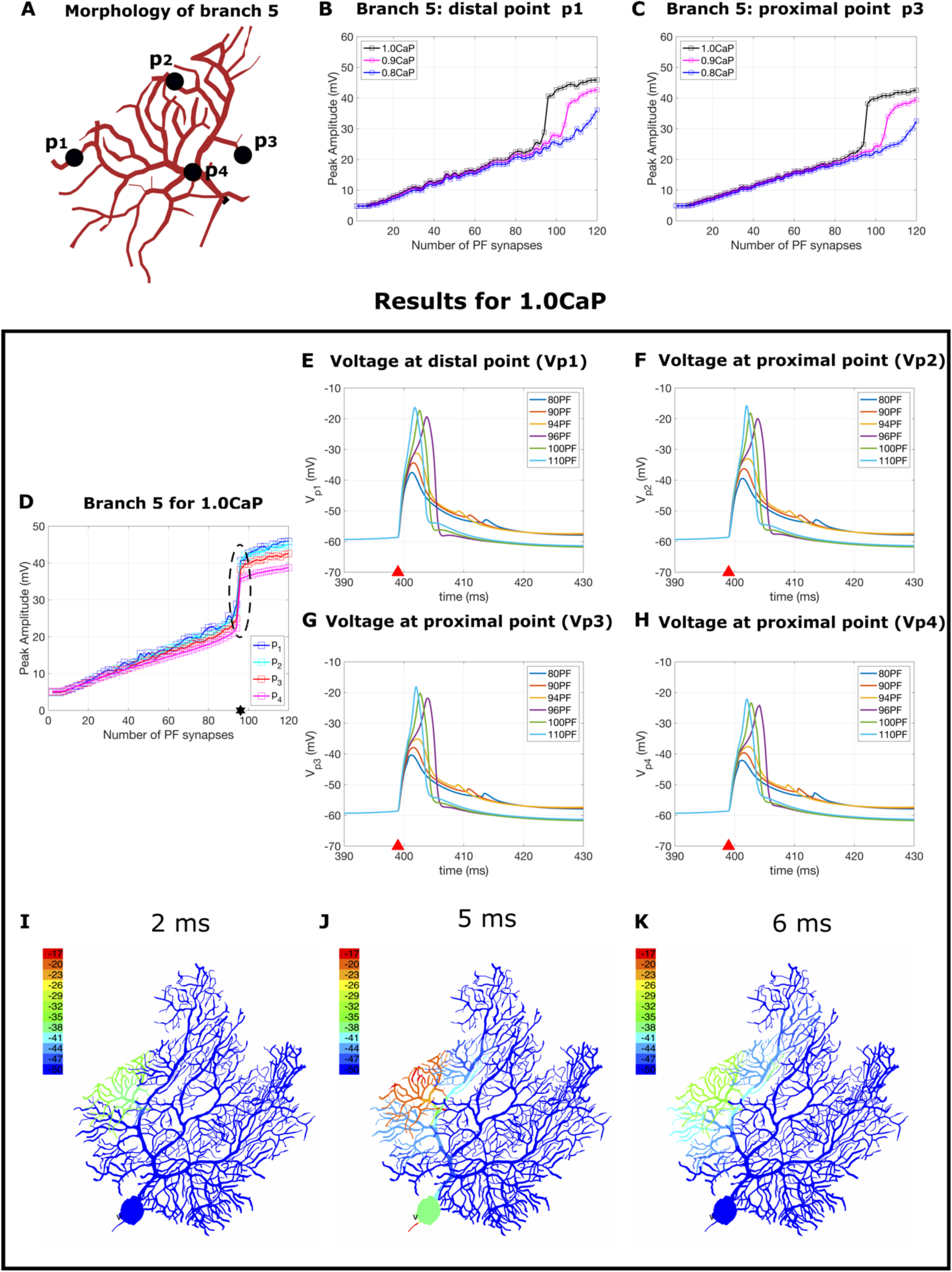
A. Illustration of branch 5. The voltage is measured at four points: two distal point *p*_1_ and *p*_2_, and two proximal points *p*_3_ and *p*_4_. B. and C. PAR for the distal point *p*_1_ and for the proximal point *p*_3_, respectively with increasing number of activated PF synapses. The black line indicates the response when considering the reference value of P-type calcium channel conductance density (CaP_g), while the magenta and blue lines show relative decreases in P-type calcium channel conductance density of 10% and 20%, respectively. D. PAR for the four points for the reference value of 1.0 CaP_g. The bimodal linear-step-plateau response (circled) occurs at a threshold of 96 PF (shown using a star). E, F, G and H. Voltage response measured at the points *p*_1_ – *p*_4_ for different number of activated PF in [10,120]. I, J and K. Dendritic spike propagation 2ms, 5ms and 6ms after activating the PF input. Observe that the dendritic spikes initiate at the tip of branch 5 and propagate towards the smooth dendrite, depolarizing the entire branch. After 5 ms from activation, the depolarization on branch 5 reaches −17mV, after which we see a slight depolarization towards the neighboring branches 3, 4, 6 and 7. The maximum depolarization detected on a nearby branch is of −41mV and occurs on branch 4.

Observe that branch 5 (**Figure S3**) attains its bimodal linear step-plateau response for its baseline value of 1.0 CaP_g and triggers a significant jump response which finishes at 96 PF (**Figure S3D**). When studying the dendritic spike propagation corresponding to 96 activated PF on branch 5, we observe that the dendritic spikes attain maximum depolarization (−17mV) after approximately 5ms (**Figure S3J**), after which they begin to spread slightly towards their neighbors: branch 4 (−41mV), branches 3, 6 and 7 (−44mV).

On the other hand, branch 10 (**Figure S4**) originally exhibited a linear response, which was converted into bimodal by increase CaP_g by 50%. The bimodal linear-step-plateau response is shown in **Figure S4D** and corresponds to a threshold of 44PF. The dendritic spikes originating in branch 10, remain perfectly constrained in this branch and achieve maximum depolarization after 3ms from activation (**Figure S4I**).

Branch 11, like branch 5 discussed earlier, already had a bimodal response with a threshold of 62PF. With the modification in Kv4_g as summarized in Figure 5E, its dendritic spikes now remain perfectly constrained and do not spread towards neighboring branches 12 and 13 (**Figure S5G-I**). However, when using with baseline value of 1.0Kv4_g, these dendritic spikes do propagate and encompass the entire branch 12 and 13 (not shown).

Branch 12 (**Figure S6**) is one of the largest and more distal branches in our PC model, it was already shown to attain a bimodal response.^27^ Here, we show the results when using a Kv4_g as in Figure 5E and **Table S2**.

We observed that decreasing its baseline CaP_g conductance by 20%, maintains its bimodal response, showing a significant jump at a PF threshold of 72PF. On the other hand, reducing its CaP_g by 50% results in a linear response (gray line in **Figure S6B**). Its dendritic spikes propagation shows how initially the spikes reach maximum depolarization on the left side of branch 12 (**Figure S6H**), after which they spread towards the right side, where due to its proximity to neighboring branch 13, they start causing significant depolarization (**Figure S6I**). Branch 13 is fully depolarized after 8ms from activation (**Figure S6J**), after which the spikes advance downwards partially depolarizing branches 9 and 10 (**Figure S6K**). Ultimately, after 10ms from activation, the spikes reach branch 11, causing a −17mV depolarization on the tip, after which the dendritic spikes subside. Observe that the final **Figure S6L** looks very similar to the activation of branch 11 (**Figure S5H**).

Without Kv4_g increases as shown in Figure 4E, branch 12 causes a much faster and complete depolarization of the entire middle division, reaching −17mV on branch 9-13. We have tried increasing Kv4_g further, but it was impossible to stop depolarization of branch 13. This is a very small branch, that requires a very large 2.2CaP_g for attaining a bimodal response and is situated in the immediate vicinity of the right side of branch 12. All our experiments showed that as soon as the dendritic spikes reach the right side of branch 12, branch 13 follows, acting intertwined, as if they were one single branch.

In conclusion, this particularly extreme branch in the dendritic tree, shows us that while our heterogenous model can very successfully compensate for morphological features, there are some affinities for propagation and some preferred functional processing units that do not correspond to our manually defined branches. Our branch selection was done as in the previous work^27^, with respect to morphological features but of course, there are countless manners in which these branches can be redefined.

**Figure S4:**
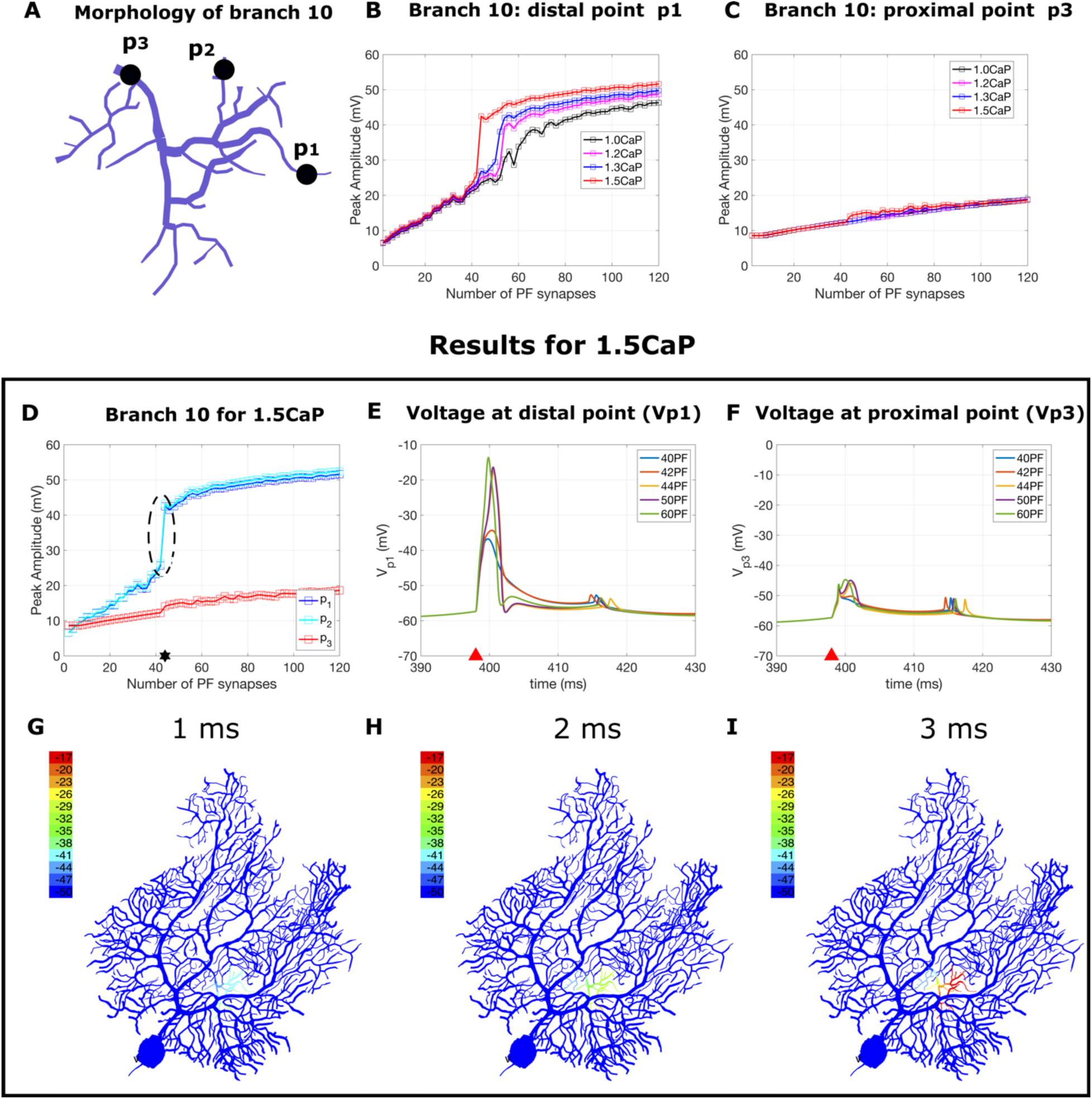
A. Illustration of branch 10. The voltage is measured at three points: two distal point *p*_1_ and *p*_2_ a proximal point *p*_3_. B and C. PAR for the distal point *p*_1_ and for the proximal point *p*_3_, respectively with increasing number of activated PF synapses. The black line indicates the response when considering the reference value of P-type calcium channel conductance density (CaP_g), while the magenta, blue and red lines show relative increases in P-type calcium channel conductance density of 20%, 30% and 50%,, respectively. D. PAR for the three points for an increase of 50% in CaP_g. The bimodal linear-step-plateau response (circled) occurs at a threshold of 44PF (shown using a star). E and F. Voltage response measured at the point *p*_1_ and *p*_3_ for different number of activated PF in [10,120] for the case of 50% increase in CaP_g. G, H and I. Dendritic spike propagation 1ms, 2ms and 3ms after activating the PF Input. Observe that the dendritic spikes initiate at the tip of branch 10 and propagate towards the smooth dendrite, depolarizing the entire branch. This depolarization is entirely localized, no spreading to neighboring branches is observed.

**Figure S5:**
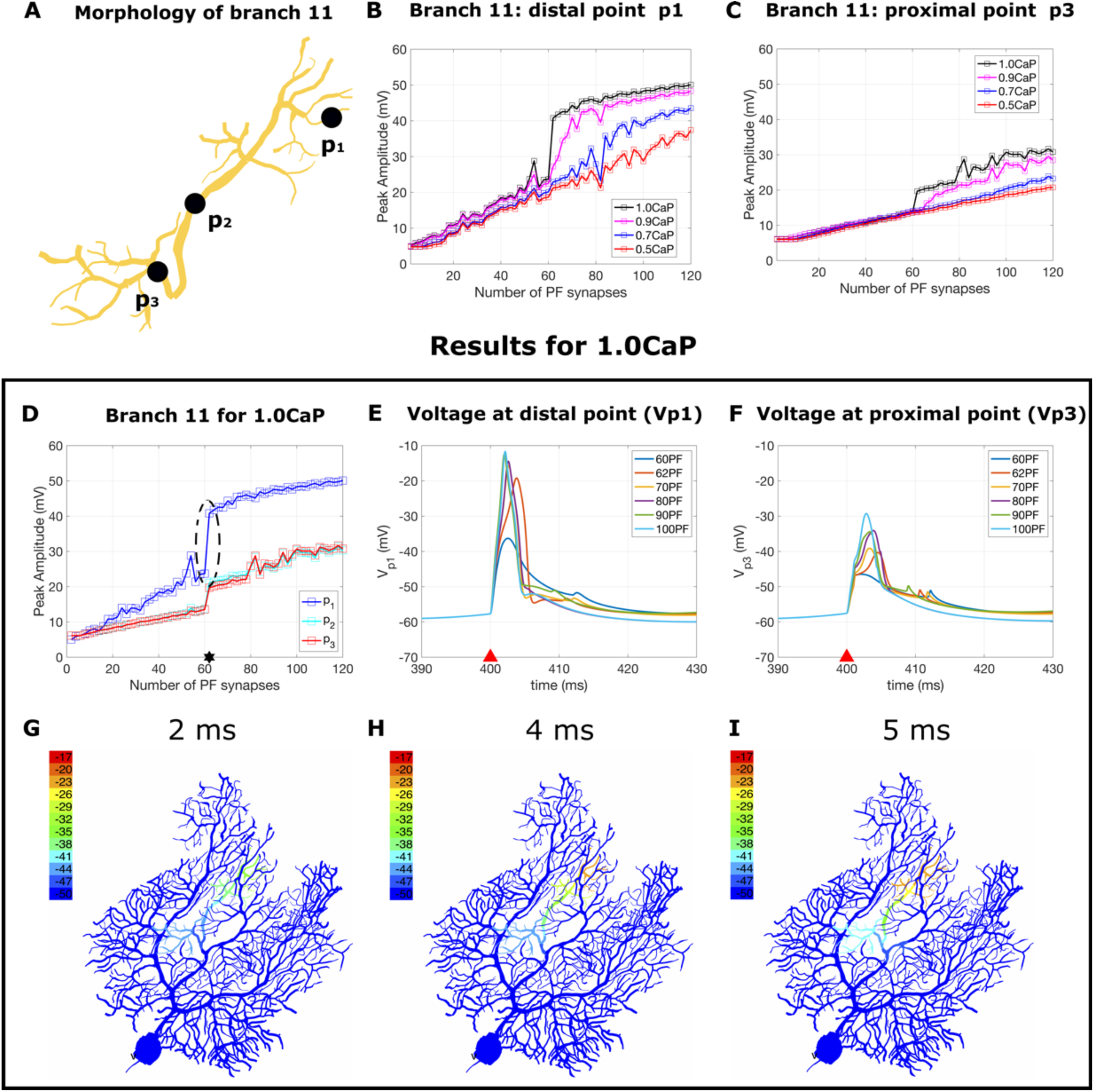
A. Illustration of branch 11. The voltage is measured at three points: a distal point *p*_1_, a middle point *p*_2_ and a proximal point *p*_3_. B and C. PAR for the distal point *p*_1_and for the proximal point *p*_3_ with increasing number of activated PF synapses. The black line indicates the response when considering the reference value of P-type calcium channel conductance density (CaP_g), while the magenta, blue and red lines show relative decreases in P-type calcium channel conductance density of 10%, 30%, respectively 50%. D. PAR for the three points for a baseline value of CaP_g. The bimodal linear-step-plateau response (circled) occurs at a threshold of 62PF (shown using a star). E and F. Voltage response measured at the point *p*_1_ and *p*_3_ for different number of activated PF in [10,120]. G, H and I. Dendritic spike propagation 2ms, 4ms and 5ms after activating the PF input. Observe that the dendritic spikes initiate at the tip of branch 11 and propagate towards the smooth dendrite, depolarizing the branch. This depolarization is entirely localized and no spreading to neighboring branches is observed.

**Figure S6:**
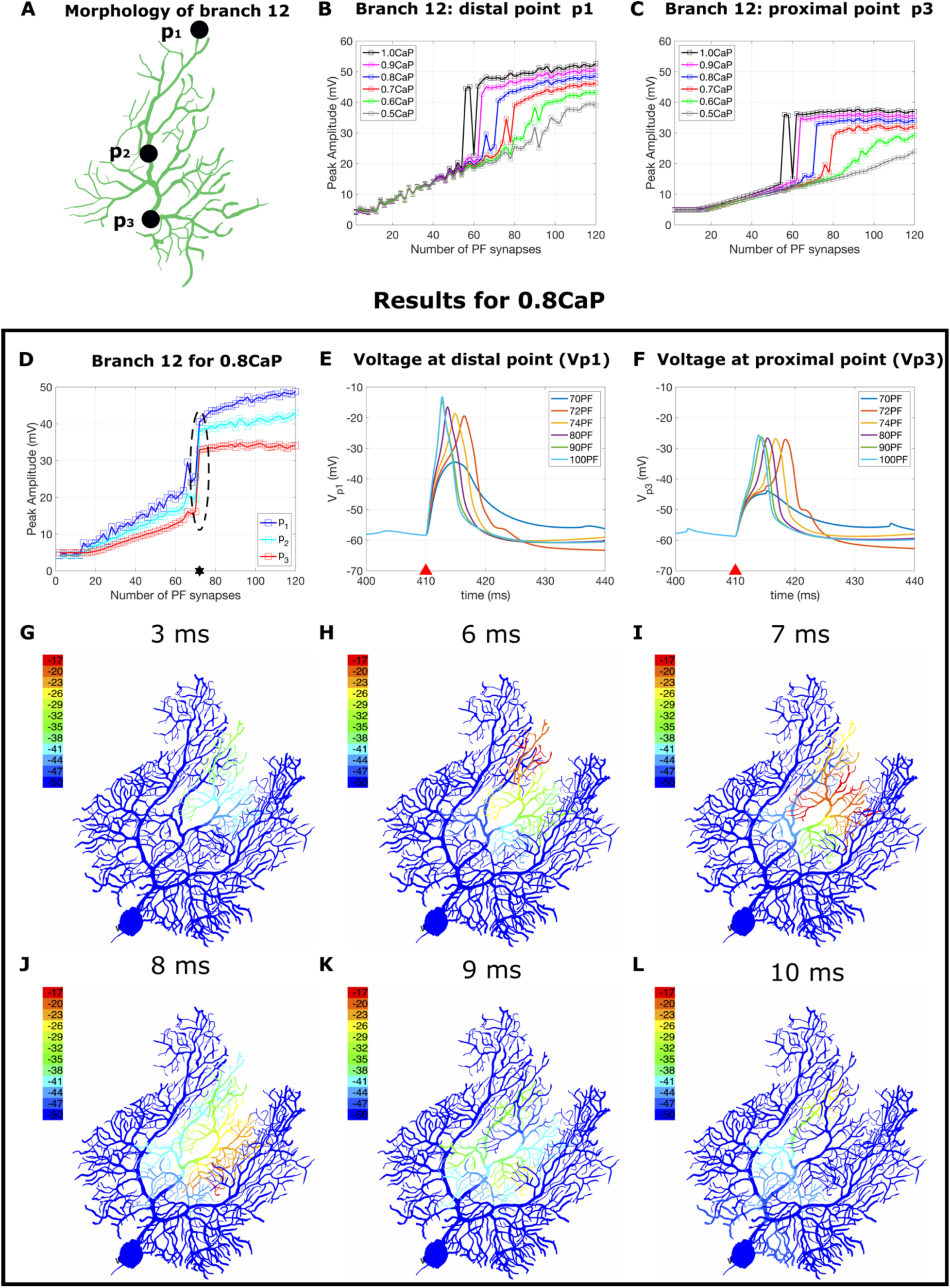
A. Illustration of branch 12. The voltage is measured at three points: a distal point *p*_1_, a middle point *p*_2_ and a proximal point *p*_3_. B. and C. PAR for the distal point *p*_1_and for the proximal point *p*_3_ with increasing number of activated PF synapses. The black line indicates the response when considering the reference value of P-type calcium channel conductance density (CaP_g), while the magenta, blue, red, green and gray lines show relative decreases in P-type calcium channel conductance density of 10%, 20%, 30%, 40% and 50%, respectively. D. PAR for the three points for a decrease of 20% in the value of CaP_g. The bimodal linear-step-plateau response (circled) occurs at a threshold of 72PF (shown using a star). E and F. Voltage response measured at the point p1 and p3 for different number of activated PF in [10,120]. G-L. Dendritic spike propagation 3ms, 6ms, 7ms, 8ms, 9ms and 10ms after activating the PF input. Observe that the dendritic spikes initiate at the tip of branch 12 and propagate towards the smooth dendrite, depolarizing the branch. First, branch 12 reaches maximum depolarization in its left side (panel H), after which it encompasses the entire branch and spreads towards neighboring small branch 13 (panel I). After 8-10ms, the depolarization within branch 12 begins subsiding and it then spreads towards branches 11, 9 and 10 (panel J-K), while continuing to fully depolarize branch 11 at t = 10ms, with the tip of branch 11 reaching −17mV.

### 4. Heterogenous model validation: climbing fiber activation and frequency-current curve

**Figure S7A** shows the results obtained when simulating climbing fiber input, we observe the resulting complex spike. **Figure S7B** shows the F-I curve for the heterogenous model (in red) compared to the homogenous model^27^ (blue).

**Figure S7:**
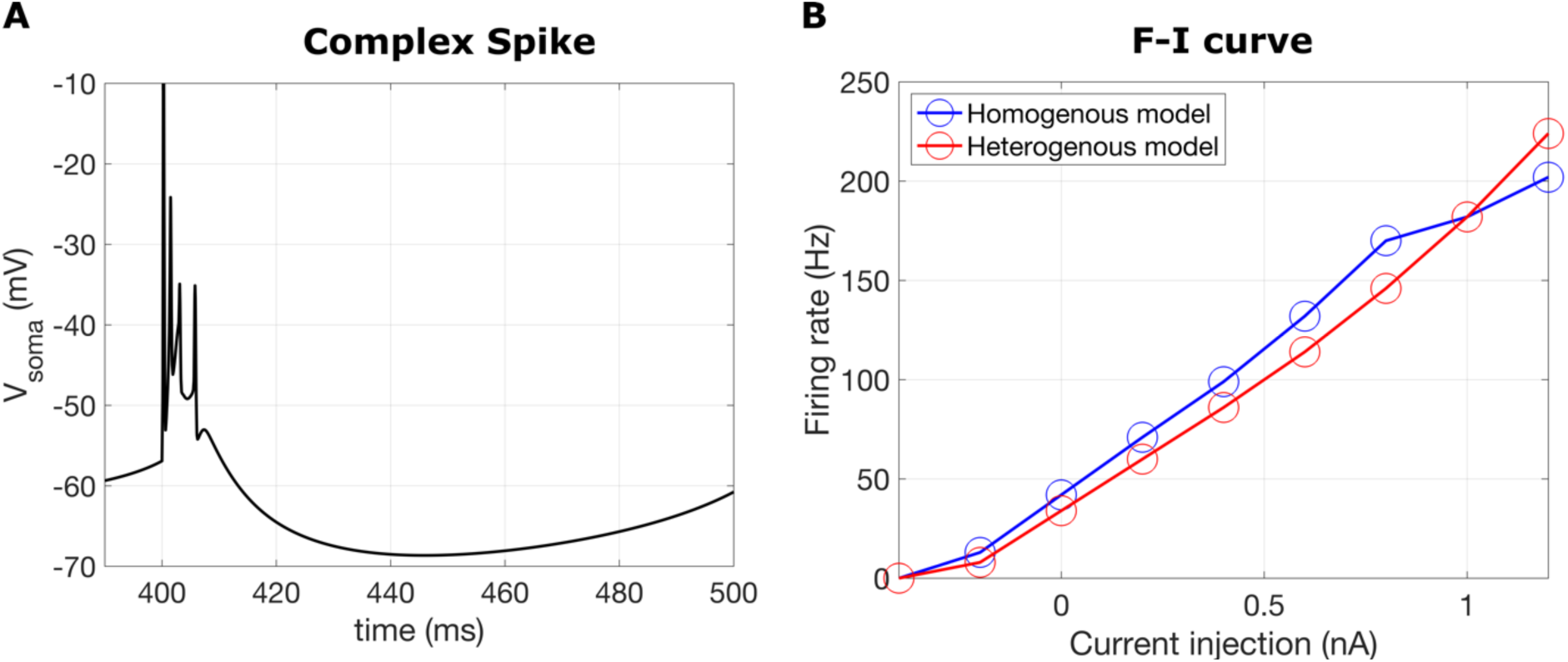
Validation of the heterogenous model. **A.** Somatic complex spike recorded with the heterogenous ion channel density model when climbing fiber input is activated at t = 400ms. **B.** F-I curve for the heterogenous model (red), compared to the homogenous model (blue).

**Table S1.** Baseline parameters^27^ for soma, axon, smooth dendrites, and spiny dendrites for the heterogenous ion channel.

| Conductance densities<br>($nS/cm^2$) | Soma | Axon | Smooth Dendrites | Spiny Dendrites |
| --- | --- | --- | --- | --- |
| Sk2 | 0.0075 | 0.0278 | $3.6 \cdot 10^{-4}$ | $3.6 \cdot 10^{-4}$ |
| Kv4 | 0 | 0 | 0.0252 | 0.0264 |
| Kv4s | 0 | 0 | 0.015 | 0.015 |
| Hpkj | $1.08 \cdot 10^{-4}$ | $1.08 \cdot 10^{-4}$ | $3.6 \cdot 10^{-4}$ | $3.24 \cdot 10^{-4}$ |
| Kv1 | 0 | 0 | 0.002 | 0.001 |
| Kv3 | 1.8 | 115.2 | 0.1512 | 0.1512 |
| CAT3 | 0 | $1.28 \cdot 10^{-4}$ | $2.7 \cdot 10^{-5}$ | $1.08 \cdot 10^{-4}$ |
| CaP | $1.9 \cdot 10^{-4}$ | 0.0023 | $1.9 \cdot 10^{-4}$ | $7.6 \cdot 10^{-4}$ |
| mslo | 0.8736 | 6 | 0.215 | 0.0448 |
| NaRsg | 0.13168 | 0.56 | 0 | 0 |
| NaP | $1.4 \cdot 10^{-4}$ | 0.0023 | 0 | 0 |
| abBK | 0.3 | 1.05 | 0 | 0.15 |

**Table S2.** Heterogenous ion channel model: maximum conductance densities of the ion channels for each branch in the PC dendritic tree. Unlike the previous model, where all branches were characterized by the same conductance densities given in the last column of **Table S1**, in the novel model each activated branch has its set of own set of conductances such that each branch attains a bimodal linear-step-plateau response. The maximum conductance densities of the ion channels corresponding to the other compartments (soma, axon, and smooth dendrites) are unchanged (see **Table S1**).

| Branch | Sk2<br>($nS/cm^2$) | Kv4<br>($nS/cm^2$) | Kv4s<br>($nS/cm^2$) | Hpkj<br>($nS/cm^2$) | Kv1<br>($nS/cm^2$) | Kv3<br>($nS/cm^2$) | CAT3<br>($nS/cm^2$) | CaP<br>($nS/cm^2$) | mslo<br>($nS/cm^2$) |
| --- | --- | --- | --- | --- | --- | --- | --- | --- | --- |
| 1 | $3.6 \cdot 10^{-4}$ | 0.0264 | 0.015 | $3.24 \cdot 10^{-4}$ | 0.001 | 0.1512 | $1.08 \cdot 10^{-4}$ | $1.1 \cdot 10^{-3}$ | 0.0896 |
| 2 | $3.6 \cdot 10^{-4}$ | 0.0264 | 0.015 | $3.24 \cdot 10^{-4}$ | 0.001 | 0.1512 | $1.08 \cdot 10^{-4}$ | $1.21 \cdot 10^{-3}$ | 0.0896 |
| 3 | $3.6 \cdot 10^{-4}$ | 0.0264 | 0.015 | $3.24 \cdot 10^{-4}$ | 0.001 | 0.1512 | $1.08 \cdot 10^{-4}$ | $1.29 \cdot 10^{-3}$ | 0.0896 |
| 4 | $3.6 \cdot 10^{-4}$ | 0.0264 | 0.015 | $3.24 \cdot 10^{-4}$ | 0.001 | 0.1512 | $1.08 \cdot 10^{-4}$ | $1.11 \cdot 10^{-3}$ | 0.0896 |
| 5 | $3.6 \cdot 10^{-4}$ | 0.066 | 0.015 | $3.24 \cdot 10^{-4}$ | 0.001 | 0.1512 | $1.08 \cdot 10^{-4}$ | $7.6 \cdot 10^{-4}$ | 0.0896 |
| 6 | $3.6 \cdot 10^{-4}$ | 0.0739 | 0.015 | $3.24 \cdot 10^{-4}$ | 0.001 | 0.1512 | $1.08 \cdot 10^{-4}$ | $1.67 \cdot 10^{-3}$ | 0.0896 |
| 7 | $3.6 \cdot 10^{-4}$ | 0.0924 | 0.015 | $3.24 \cdot 10^{-4}$ | 0.001 | 0.1512 | $1.08 \cdot 10^{-4}$ | $2.13 \cdot 10^{-3}$ | 0.0896 |
| 8 | $3.6 \cdot 10^{-4}$ | 0.0792 | 0.015 | $3.24 \cdot 10^{-4}$ | 0.001 | 0.1512 | $1.08 \cdot 10^{-4}$ | $6.84 \cdot 10^{-4}$ | 0.0896 |
| 9 | $3.6 \cdot 10^{-4}$ | 0.0264 | 0.015 | $3.24 \cdot 10^{-4}$ | 0.001 | 0.1512 | $1.08 \cdot 10^{-4}$ | $9.12 \cdot 10^{-4}$ | 0.0896 |
| 10 | $3.6 \cdot 10^{-4}$ | 0.0264 | 0.015 | $3.24 \cdot 10^{-4}$ | 0.001 | 0.1512 | $1.08 \cdot 10^{-4}$ | $1.11 \cdot 10^{-3}$ | 0.0896 |
| 11 | $3.6 \cdot 10^{-4}$ | 0.0396 | 0.015 | $3.24 \cdot 10^{-4}$ | 0.001 | 0.1512 | $1.08 \cdot 10^{-4}$ | $7.6 \cdot 10^{-4}$ | 0.0896 |
| 12 | $3.6 \cdot 10^{-4}$ | 0.0396 | 0.015 | $3.24 \cdot 10^{-4}$ | 0.001 | 0.1512 | $1.08 \cdot 10^{-4}$ | $6.08 \cdot 10^{-4}$ | 0.0896 |
| 13 | $3.6 \cdot 10^{-4}$ | 0.066 | 0.015 | $3.24 \cdot 10^{-4}$ | 0.001 | 0.1512 | $1.08 \cdot 10^{-4}$ | $1.67 \cdot 10^{-3}$ | 0.0896 |
| 14 | $3.6 \cdot 10^{-4}$ | 0.0264 | 0.015 | $3.24 \cdot 10^{-4}$ | 0.001 | 0.1512 | $1.08 \cdot 10^{-4}$ | $1.21 \cdot 10^{-3}$ | 0.0896 |
| 15 | $3.6 \cdot 10^{-4}$ | 0.0264 | 0.015 | $3.24 \cdot 10^{-4}$ | 0.001 | 0.1512 | $1.08 \cdot 10^{-4}$ | $6.08 \cdot 10^{-4}$ | 0.0896 |
| 16 | $3.6 \cdot 10^{-4}$ | 0.0264 | 0.015 | $3.24 \cdot 10^{-4}$ | 0.001 | 0.1512 | $1.08 \cdot 10^{-4}$ | $9.12 \cdot 10^{-4}$ | 0.0896 |
| 17 | $3.6 \cdot 10^{-4}$ | 0.0264 | 0.015 | $3.24 \cdot 10^{-4}$ | 0.001 | 0.1512 | $1.08 \cdot 10^{-4}$ | $8.36 \cdot 10^{-4}$ | 0.0896 |
| 18 | $3.6 \cdot 10^{-4}$ | 0.0264 | 0.015 | $3.24 \cdot 10^{-4}$ | 0.001 | 0.1512 | $1.08 \cdot 10^{-4}$ | $2.13 \cdot 10^{-3}$ | 0.0896 |
| 19 | $3.6 \cdot 10^{-4}$ | 0.0264 | 0.015 | $3.24 \cdot 10^{-4}$ | 0.001 | 0.1512 | $1.08 \cdot 10^{-4}$ | $19.8 \cdot 10^{-3}$ | 0.0896 |
| 20 | $3.6 \cdot 10^{-4}$ | 0.0264 | 0.015 | $3.24 \cdot 10^{-4}$ | 0.001 | 0.1512 | $1.08 \cdot 10^{-4}$ | $2.3 \cdot 10^{-3}$ | 0.0896 |
| 21 | $3.6 \cdot 10^{-4}$ | 0.0264 | 0.015 | $3.24 \cdot 10^{-4}$ | 0.001 | 0.1512 | $1.08 \cdot 10^{-4}$ | $6.08 \cdot 10^{-4}$ | 0.0896 |
| 22 | $3.6 \cdot 10^{-4}$ | 0.0264 | 0.015 | $3.24 \cdot 10^{-4}$ | 0.001 | 0.1512 | $1.08 \cdot 10^{-4}$ | $6.08 \cdot 10^{-4}$ | 0.0896 |

## Notes

### Competing Interest Statement

The authors have declared no competing interest.

